# Intrinsically disordered regions facilitate Msn2 target search to drive promoter selectivity

**DOI:** 10.1101/2025.05.23.654710

**Authors:** Nir Strugo, Carmit Burstein, Sk Saddam Hossain, Noam Nago, Moshe Goldsmith, Hadeel Khamis, Ariel Kaplan

## Abstract

Transcription factors (TFs) regulate gene expression by binding specific DNA motifs, yet only a fraction of putative sites is occupied *in vivo*. Intrinsically disordered regions (IDRs) have emerged as key contributors to promoter selectivity, but the underlying mechanisms remain incompletely understood. Here, we use single-molecule optical tweezers to dissect how IDRs influence DNA binding by Msn2, a yeast stress-response regulator. We show that IDRs facilitate initial non-specific association with DNA and promote one-dimensional diffusion toward target motifs, supported by charge-mediated interactions. Remarkably, the IDR-dependent search mechanism displays sequence sensitivity, with promoter-derived sequences enhancing both initial binding and sliding rates, demonstrating that Msn2–DNA interactions alone are sufficient to confer promoter selectivity in the absence of chromatin or cofactors. These findings provide direct mechanistic evidence for how IDRs tune transcription factor search dynamics and expand sequence recognition beyond canonical motifs, supporting a mechanism for promoter selectivity in complex genomic contexts.

## INTRODUCTION

Eukaryotic transcription factors (TFs) orchestrate gene expression by binding to specific DNA motifs in promoters and enhancers, yet only a subset of these short and degenerate motifs present in the genome is occupied *in vivo*^1,2^. Recent work has highlighted a critical role for intrinsically disordered regions (IDRs) in conferring promoter specificity, showing that many yeast TFs, including zinc-finger, bZIP, and homeodomain proteins, require extensive IDRs for selective promoter binding^3,4^. This IDR-mediated selectivity implies an additional layer of specificity beyond the canonical DNA-binding domain (DBD), but the molecular mechanisms underlying the IDRs’ role remain elusive.

Characterized by their lack of stable secondary or tertiary structure, IDRs exhibit high conformational flexibility and multivalency, which allow them to participate in dynamic and context-dependent interactions. IDRs have been implicated in diverse regulatory functions *in vivo*: they drive transcriptional activation through recruitment of coactivators, assemble phase-separated condensates that concentrate transcriptional machinery, and modulate chromatin residence times^5,6^. In mammalian cells, disordered regions of TFs such as p53, NFκB, and the glucocorticoid receptor, contribute to spatial confinement or prolonged chromatin engagement^7–9^. However, the direct contributions of IDRs to TF–DNA interactions and promoter selectivity remain untested.

TF–DNA binding is governed not only by binding affinity, but also by the kinetics of how sites are located and engaged^10^. According to the facilitated diffusion model, TFs alternate between three-dimensional diffusion in solution and one-dimensional diffusion along the DNA, reducing search dimensionality and enhancing motif binding rates^11^. This mechanism allows efficient scanning of genomic DNA, while discriminating between cognate and non-cognate sites through transient contacts, dwell-time heterogeneity, and multistate binding behaviors^12–15^. Recent single-molecule studies have further revealed that sequence specificity can arise primarily from differences in association rates rather than dissociation kinetics, and that TFs can explore broad DNA surfaces and bypass potential targets during search^16,17^. IDRs may modulate these kinetic processes by promoting initial electrostatic engagement, stabilizing transient complexes, or tuning one-dimensional diffusion dynamics^18,19^. In parallel, several studies have shown that TF binding *in vivo* is influenced by DNA sequence features outside the core motif, such as flanking sequence composition, DNA shape, or chromatin architecture^20–22^. These observations raise the possibility that disordered regions enable TFs to sense contextual features and bias their search toward biologically relevant sites. However, the direct contributions of IDRs to these kinetic parameters, i.e. association rates, dwell-time stability, and sliding behavior, remain untested at the single-molecule level.

To directly address these questions, we use single-molecule optical tweezers to investigate how IDRs influence DNA binding by Msn2, a master regulator of the environmental stress response in budding yeast^23^. *S. cerevisiae* Msn2, composed of 704 amino acids, contains a 62-residue canonical DBD that targets stress-responsive promoter elements, flanked by extensive intrinsically disordered regions^24^ (Figure 1A) required for its *in vivo* promoter specificity and regulatory function^25^. Building on our previous single-molecule studies of transcription factor binding dynamics^26,27^, we developed assays to probe specific interactions (binding at the cognate recognition motif) and non-specific interactions (association with other DNA regions), as well as Msn2’s ability to undergo one-dimensional (1D) diffusion along DNA. These experiments allow us to dissect how disordered regions contribute to promoter-selective binding and to determine their specific roles in TF–DNA recognition.

**Figure 1:**
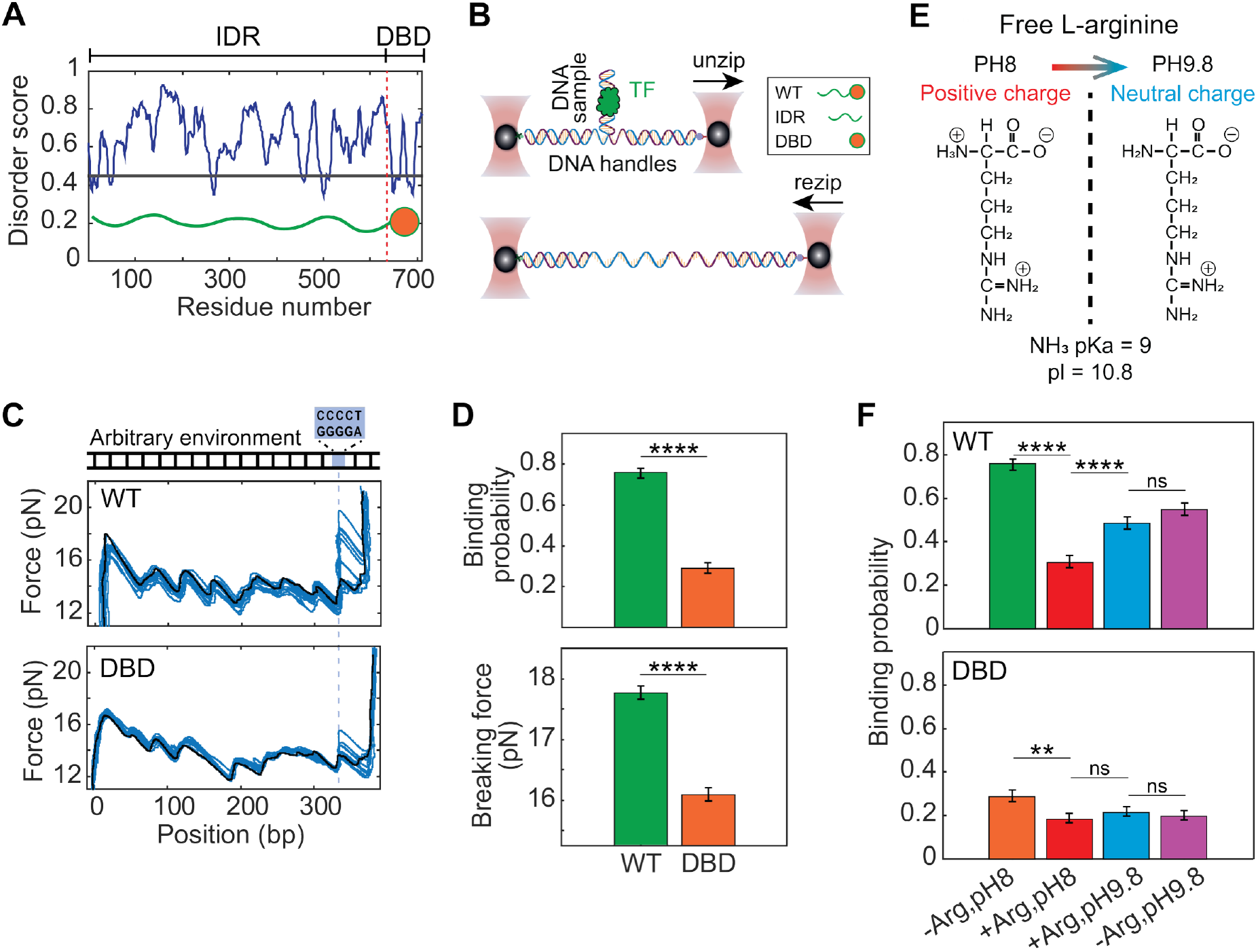
IDRs increase Msn2 binding affinity via charge-mediated interactions. (A) Disorder prediction profile of Msn2 calculated using IUPred3, highlighting the intrinsically disordered regions (IDRs) and DNA-binding domain (DBD). (B) Schematic representation of the DNA unzipping-rezipping experimental assay used to compare binding properties of wild-type Msn2 (WT), IDR-only (IDR), and DBD-only (DBD) variants. See also Figure S1. (C) Representative force-position traces from DNA unzipping experiments in the presence of 50 nM WT (top) and DBD (bottom) variants. The DNA construct contains an AT-rich sequence flanking the binding motif (position marked by dashed line). The black trace shows unzipping in a protein-free solution for reference. (D) Binding probability (total unzipping iterations n_WT_=298, n_DBD_=289) and breaking force (n_WT_=226, n_DBD_=83) for WT and DBD variants. Data shown as mean ± SEM, ****P<0.0001, χ^2^ test and Student’s t-test, respectively. (E) Charge state of free L-arginine at pH 8.0 (left, positively charged) and pH 9.8 (right, neutral) illustrating the pH-dependent protonation of the amino group (pKa=9.0). (F) Binding probability under different electrostatic perturbation conditions for WT (top panel; total unzipping iterations n_-Arg,pH8_=298, n_+Arg,pH8_=258, n_+Arg,pH9.8_=319, n_-Arg,pH9.8_=302) and DBD variants (bottom panel; n_-Arg,pH8_=289, n_+Arg,pH8_=309, n_+Arg,pH9.8_=339, n_-Arg,pH9.8_=342). Conditions from left to right: “unperturbed” (similar to D), electrostatic screening with positively charged L-arginine at pH 8.0, neutralized screening with L-arginine at pH 9.8, and control at pH 9.8 without L-arginine. Data shown as mean ± SEM, **P<0.01, ****P<0.0001, χ^2^ test. See also Table S5.

## RESULTS

### IDRs increase Msn2 binding affinity via charge-mediated interactions

To identify the contribution of IDRs to DNA binding, we compared the binding performance of wild-type Msn2 (WT) and an IDR-truncated mutant (DBD) using DNA unzipping with optical tweezers (Figure 1B; see also Figure S1 and Table S1). In previous work^26^, we showed that a bound TF hinders the propagation of the DNA unzipping fork, allowing us to measure the protein’s position and breaking force. Repeated measurements, where the system is allowed to thermally equilibrate between iterations, reveal the binding probability, a direct measure of the TF’s affinity for its binding site. In initial experiments, we used a section of the *Hor7* gene promoter, which contains four AGGGG binding motifs (Figure S2A) and is known to be bound by Msn2 *in vivo*^25^. However, the relatively small force signature of Msn2, which possesses only two zinc-fingers, combined with the high-force background from the flanking sequences, hindered the identification of binding events, and we were able to observe events only at one of the four sites (Figure S2A). To improve sensitivity, we designed a dedicated DNA construct with a single Msn2 binding motif, flanked by a short AT-rich sequence (which requires lower unzipping forces) and preceded by a ∼300 bp segment lacking binding motifs. This design minimized background forces and enabled clearer detection of binding events (Figure 1C). Repeated unzipping experiments with WT and DBD revealed a significant decrease in both binding probability and breaking force upon IDR removal (Figure 1D), indicating that Msn2’s IDRs enhance its binding affinity for the binding motif. Notably, we also tested a variant containing only the IDRs (IDR; Figure S1 and Table S1) and observed no detectable unzipping signature at the motif.

Next, we sought to understand the mechanism by which IDRs increase the affinity of Msn2 for DNA. We hypothesized that charge-mediated interactions might contribute and modulated them by introducing 50 mM free L-arginine into the buffer (which contains 150 mM KCl). At pH 8.0, L-arginine is positively charged due to protonation of its guanidino and amino groups, while its carboxyl group is deprotonated (Figure 1E; Tables S2,3). This treatment led to a marked reduction in binding probability for WT Msn2 (Figure 1F, top). A similar effect was observed upon increasing ionic strength with KCl alone (Figure S2B), supporting a common electrostatic screening mechanism. However, since L-arginine can also engage directly with DNA or protein surfaces, alter solvation and affect IDR conformational preferences, we further tested whether the effect depends specifically on its net charge. Raising the pH to 9.8, which neutralizes the amino group of L-arginine (pKa = 9.0) without affecting the side chains in Msn2 (Tables S2, S3), partially restored binding. Notably, at pH 9.8, the binding probability was similar regardless of whether L-arginine was present, further supporting the conclusion that the inhibition observed at pH 8.0 reflects electrostatic screening by the charged form of L-arginine, rather than any other molecular interaction. While we cannot fully exclude additional contributions from L-arginine’s chemical properties, the observed pH sensitivity and ionic strength dependence argue that such contributions are minimal at the concentration used. The DBD variant showed a qualitatively similar trend (Figure 1F, bottom; Figure S2C), consistent with the known electrostatic interactions between zinc fingers and DNA, but the effect was substantially milder. Hence, our results suggest that the IDRs enhance DNA binding largely through charge-mediated interactions. Moreover, they establish L-arginine as a useful experimental probe for selectively perturbing IDR–DNA interactions, enabling further dissection of the role of disordered regions in transcription factors’ function.

### IDRs enhance the association rate of Msn2 to its binding motif

While our unzipping experiments demonstrated that IDRs increase Msn2’s overall binding affinity, they did not directly resolve whether this enhancement reflects changes in association rate (k_on_), dissociation rate (k_off_), or both. To dissect the specific kinetic parameters affected by IDRs, we exploited a method we previously developed^27^, based on monitoring thermal breathing fluctuations of DNA (Figure 2A). The DNA construct is unzipped until reaching several base-pairs upstream of the binding motif, after which the optical traps are held at a constant position. The DNA then undergoes rapid thermal fluctuations on the millisecond timescale, during which approximately 20 bp, including the binding site, transiently and repeatedly separate into single strands (“open”) and re-anneal (“close”). TF binding traps the DNA in the closed state, thereby suppressing these fluctuations and allowing to identify the bound complex. By analyzing the durations of unbound and bound states (Figure 2A), we directly measure the association and dissociation rates, respectively. Using this approach, we found that while the dissociation rate was unaffected by IDR truncation, the association rate decreased approximately sixfold (Figure 2B), indicating that IDRs enhance binding affinity by accelerating target association.

**Figure 2:**
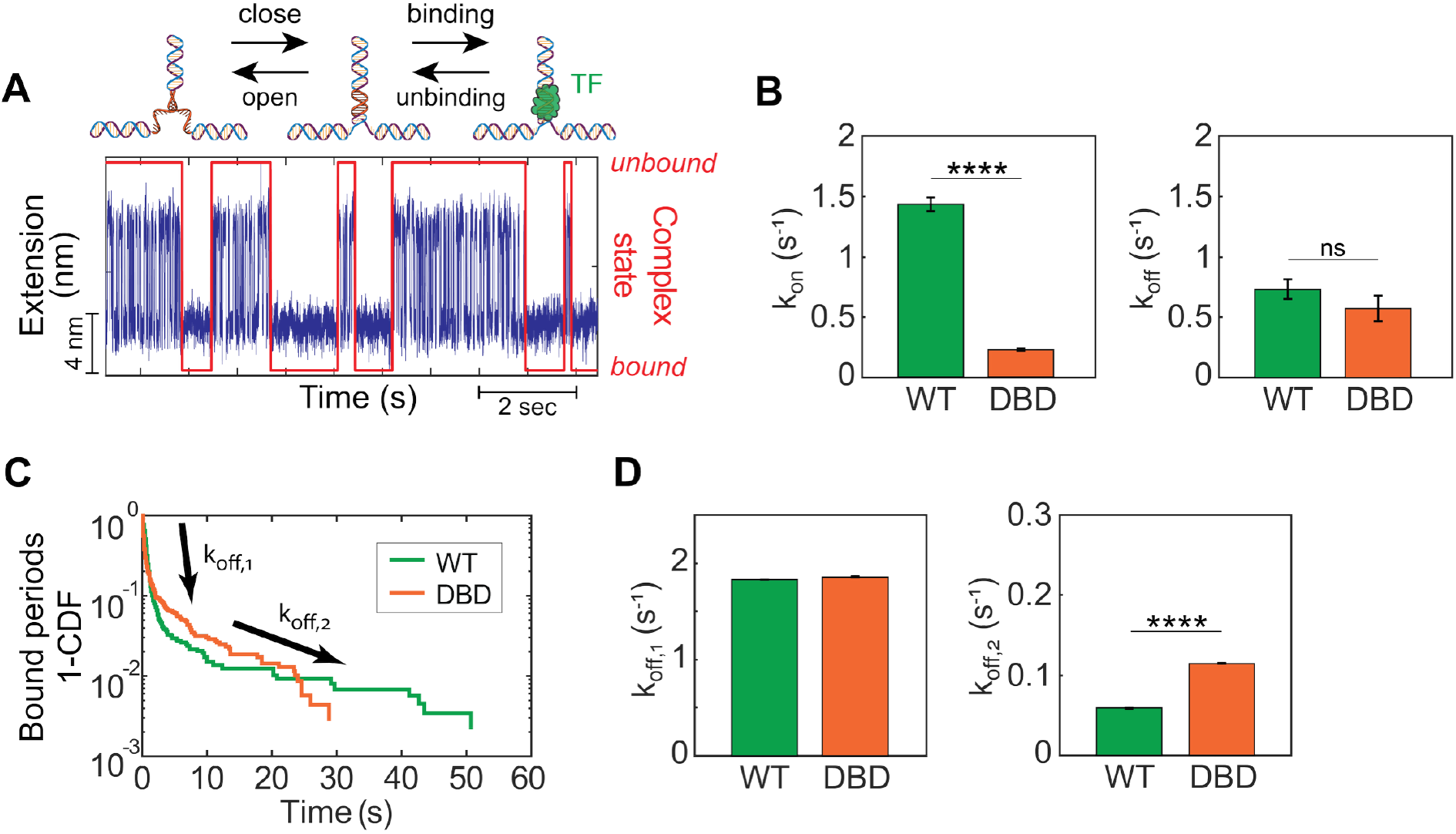
IDRs enhance the association rate of Msn2 to its binding motif. (A) Schematic illustration of the kinetic fluctuation suppression assay. After partial unzipping, the DNA construct undergoes rapid thermal fluctuations between “open” and “closed” states. When a transcription factor binds to its recognition site within the fluctuating segment, these fluctuations are suppressed. (B) Comparison of association rates (left panel; total period events n_WT_=838, n_DBD_=642) and dissociation rates (right panel; n_WT_=892, n_DBD_=702) for Msn2 WT and DBD variants at the specific binding site. Data shown as mean ± SEM, ****P<0.0001, permutation test with 10,000 resampling. (C) Cumulative distribution function (CDF) of bound state durations for both WT and DBD variants, revealing two distinct populations with different dissociation rates, k_off,1_ and k_off,2_. (D) Quantitative comparison of the two dissociation rates between WT and DBD variants, calculated from a double-exponential fit to the bound-state duration distributions (total period events n_WT_=623, n_DBD_=307). Data shown as mean ± SEM, ****P<0.0001, permutation test with 10,000 resampling. See also Table S6.

Analysis of the full probability distributions revealed further complexity. While unbound-state durations followed a single-exponential distribution (Figure S3A), the bound-state durations deviated from this, suggesting the presence of two kinetically distinct populations of Msn2–DNA bound complexes (Figure 2C). This biphasic behavior was observed for both WT and DBD variants and appears to be specific to Msn2, as it was not observed for the previously studied Egr-1 DBD (Figure S3B).

Notably, while the rapidly dissociating subpopulation showed similar lifetimes for WT and DBD, the more stable subpopulation was longer-lived for the WT protein (Figure 2D). These findings suggest that, in addition to promoting faster target association, the IDRs also stabilize a subset of tightly bound Msn2–DNA bound complexes.

### IDRs promote cooperative non-specific DNA binding by Msn2

During our analysis of the equilibrium binding experiments (Figure 1), we discovered an unexpected phenomenon that provided further insight into IDRs’ function. In some unzipping cycles, we observed binding events tens to hundreds of base pairs away from the canonical binding site (Figure 3A, S4A). These non-specific interactions were predominantly detected with WT Msn2, rarely with the DBD variant, and never in the absence of protein, indicating that the IDRs are critical for their formation (Figure 3B). However, unzipping experiments with the IDR variant showed no non-specific (nor specific) events.

**Figure 3:**
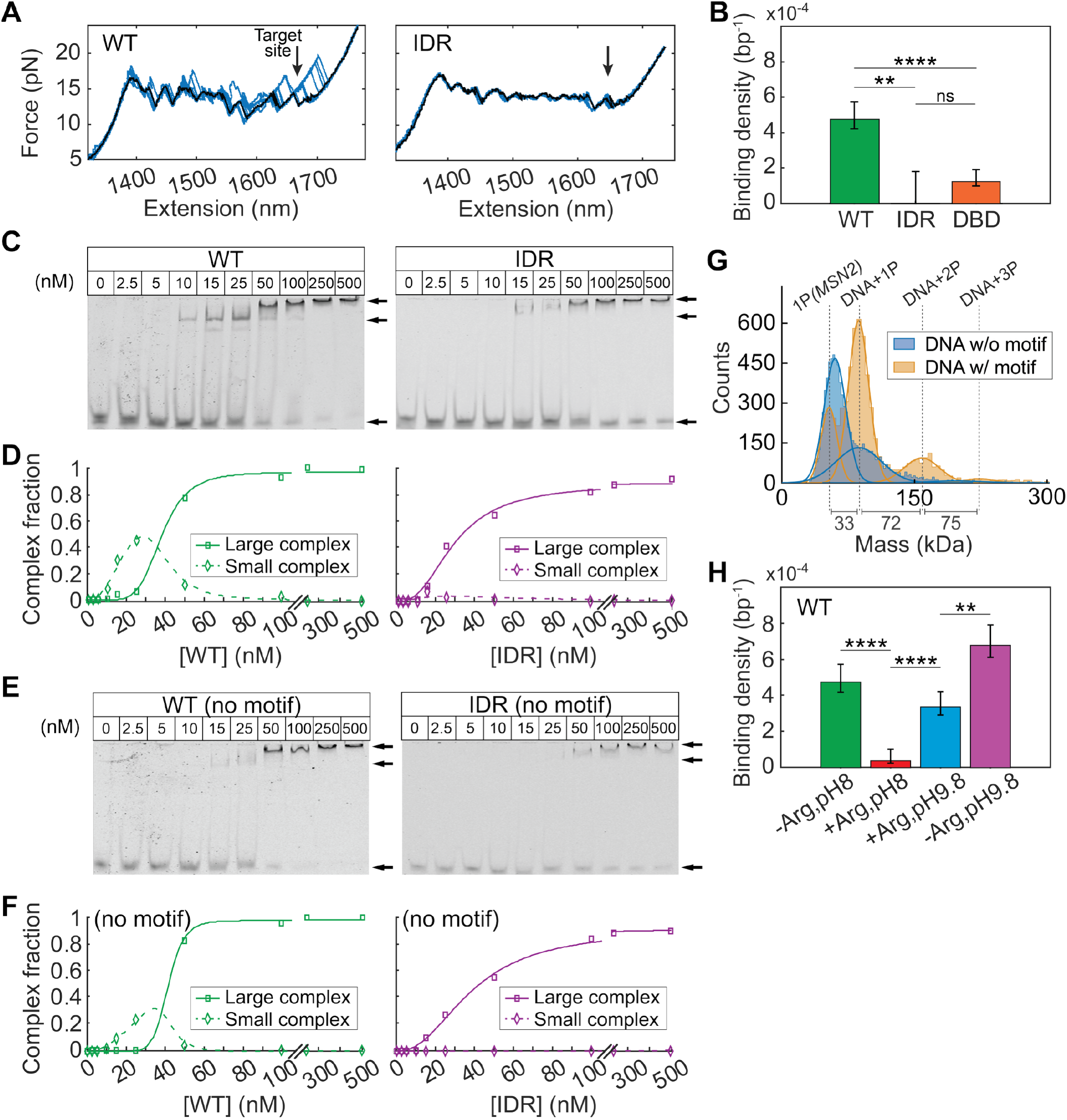
IDRs promote cooperative non-specific DNA binding by Msn2. (A) Representative force-extension traces from DNA unzipping experiments with Msn2 variants. The black trace was obtained in a protein-free solution for reference. Traces with WT Msn2 show non-specific binding events distant from the canonical binding site (marked by an arrow). (B) Non-specific binding density along the DNA construct prior to the binding site region, comparing WT and IDR variants (total position bins n_WT_=9238, n_IDR_=868, n_DBD_=8959). Data shown as mean ± SEM, **P<0.01, ****P<0.0001, χ^2^ test. (C) EMSA using fluorescently labeled 51-bp DNA fragments containing the Msn2 binding motif, with increasing concentrations of Msn2 variants. In addition to the unbound DNA band at the bottom of the gel, two distinct bound complexes are observed in the upper bands (indicated by arrows). (D) Quantitative analysis of bound complex formation as a function of protein concentration for both complexes observed in the EMSA results above. (E) EMSA with DNA sequences lacking the binding motif. (F) Quantitative analysis of complex formation for bound states observed in the motif-free EMSA. (G) Mass distribution profiles obtained by mass photometry of solutions containing WT Msn2 with 51-bp DNA fragments either containing (orange) or lacking (blue) the binding motif. Gaussian fits reveal distinct population peaks. Mass differences between adjacent peaks and the corresponding molecular identities are indicated. (H) Non-specific binding density of the WT variant under various electrostatic perturbation conditions as described in Figure 1 (total position bins n_-Arg,pH8_=9238, n_+Arg,pH8_=8401, n_+Arg,pH9.8_=9424, n_-Arg,pH9.8_=9362). Data shown as mean ± SEM, **P<0.01, ****P<0.0001, χ^2^ test. See also Table S7.

To further characterize the nature of the complexes involved in the observed non-specific binding, we conducted electrophoretic mobility shift assays (EMSA) with a 51 bp DNA segment containing the canonical binding motif. For WT Msn2, we observed two distinct bands, indicating the formation of two nucleoprotein complexes with different mobilities (Figure 3C, left). Quantification with a sequential binding model revealed that the slower-migrating, larger complex accumulates cooperatively at the expense of the smaller one (Figure 3D, left; Table S4), consistent with cooperative multimer formation. While no binding was detected for the IDR-only variant in our optical tweezers assay (Figure 3A), EMSA revealed that it does engage DNA, forming only the higher-order complex and failing to produce the smaller one observed with WT (Figure 3C, right; Figure 3D, right). Since unzipping detects proteins that interfere with strand separation, this indicates that although the IDRs contribute to non-specific binding, they either do not bridge between the strands or do so weakly enough to prevent detection by unzipping fork blockage. Together, these findings indicate that the IDRs mediate non-specific DNA interactions, but the DBD is required to stabilize these interactions and produce the unzipping signature.

To test whether these complexes require the presence of the canonical binding motif, we replaced it with a random sequence (Figure 3E). Removal of the motif completely abolished complex formation by the DBD variant (Figure S4B), whereas both WT and IDR were still able to form complexes. However, the substitution reduced the abundance of the small complex in WT while leaving the larger complex largely unchanged (Figure 3E–F). These results confirm that the smaller complex depends on the DBD and reflects both specific and non-specific interactions. In contrast, the larger complex (IDR-dependent and DBD-stabilized) reflects cooperative non-specific interactions only. Interestingly, the DBD-only variant, which showed no binding in the absence of the motif also produced two shifted bands in its presence (Figure S4B), suggesting that even motif-specific DBD– DNA interactions can generate multiple conformational or stoichiometric states. These may reflect distinct protein–DNA binding modes or DBD dimerization at the motif.

To clarify the molecular composition of the observed complexes, we used mass photometry with 50 nM WT Msn2 and measured the mass distribution of Msn2-DNA complexes with and without the binding motif (Figure 3G). The data revealed discrete peaks with consistent mass differences: a difference of ∼33 kDa between the first and second peaks (similar to the mass of the 51 bp DNA segment used in this experiment) and ∼72-75 kDa between successive peaks (similar to the mass of Msn2).

These stepwise mass differences allowed us to confidently assign the peaks as free protein (1P), DNA with one protein (DNA+1P), DNA with two proteins (DNA+2P), and DNA with three proteins (DNA+3P), respectively.

These results demonstrate that the larger complexes observed in our EMSA experiments represent discrete assemblies of multiple Msn2 molecules on individual DNA fragments, rather than non-specific aggregation. DNA containing the binding motif (orange) exhibited a prominent peak at the DNA+1P position, consistent with specific binding to the canonical motif. In contrast, DNA lacking the motif (blue) showed a more uniform distribution across multiple bound states, and a higher free-protein peak.

To test whether the formation of these complexes depends on charge-mediated interactions, we measured the density of non-specific Msn2 binding events observed in unzipping experiments under electrostatic perturbation with L-arginine. For WT Msn2, this treatment led to a clear reduction in non-specific binding (Figure 3H), which was reversed by pH increase, mirroring the behavior observed for site-specific interactions. This suggests that both binding modes rely on similar charge-mediated mechanisms. While less frequent overall, non-specific binding by the DBD-only variant was also reduced by these perturbations (Figure S4C). Taken together, the EMSA, mass photometry, and perturbation results show that IDRs support multiple, cooperative, non-specific contacts with DNA through charge-mediated interactions.

### IDRs mediate Msn2 binding to single-stranded DNA

During inspection of the rezipping traces from our equilibrium unzipping experiments, we unexpectedly observed instances of hysteresis, i.e. instances where the force during rezipping differed from the unzipping force at the same position (Figures 4A, S5A). These events, which occurred both near and far from the binding motif, reflect hindrance of DNA re-annealing and were rarely observed in the absence of protein, suggesting that Msn2 may bind to single-stranded DNA (ssDNA) exposed during unzipping. Hysteresis was significantly more pronounced with full-length Msn2 compared to the DBD and IDR variants (Figure 4B), and complementary EMSA experiments showed formation of ssDNA-bound complexes for WT and IDR, but not when the IDRs were removed (Figure 4C), suggesting that ssDNA binding is mediated primarily by the IDRs and might be stabilized by the DBD. As with double-stranded DNA (dsDNA) binding, ssDNA interactions were sensitive to L-arginine perturbation (Figure 4D, S5B). However, unlike dsDNA binding, ssDNA interactions were not rescued by raising the pH to 9.8 in the presence of L-arginine, and remained low under all high-pH conditions, including in the absence of L-arginine. This suggests that IDR–ssDNA interactions rely on a distinct mechanism, sensitive to L-arginine but not governed solely by classical electrostatic screening.

**Figure 4:**
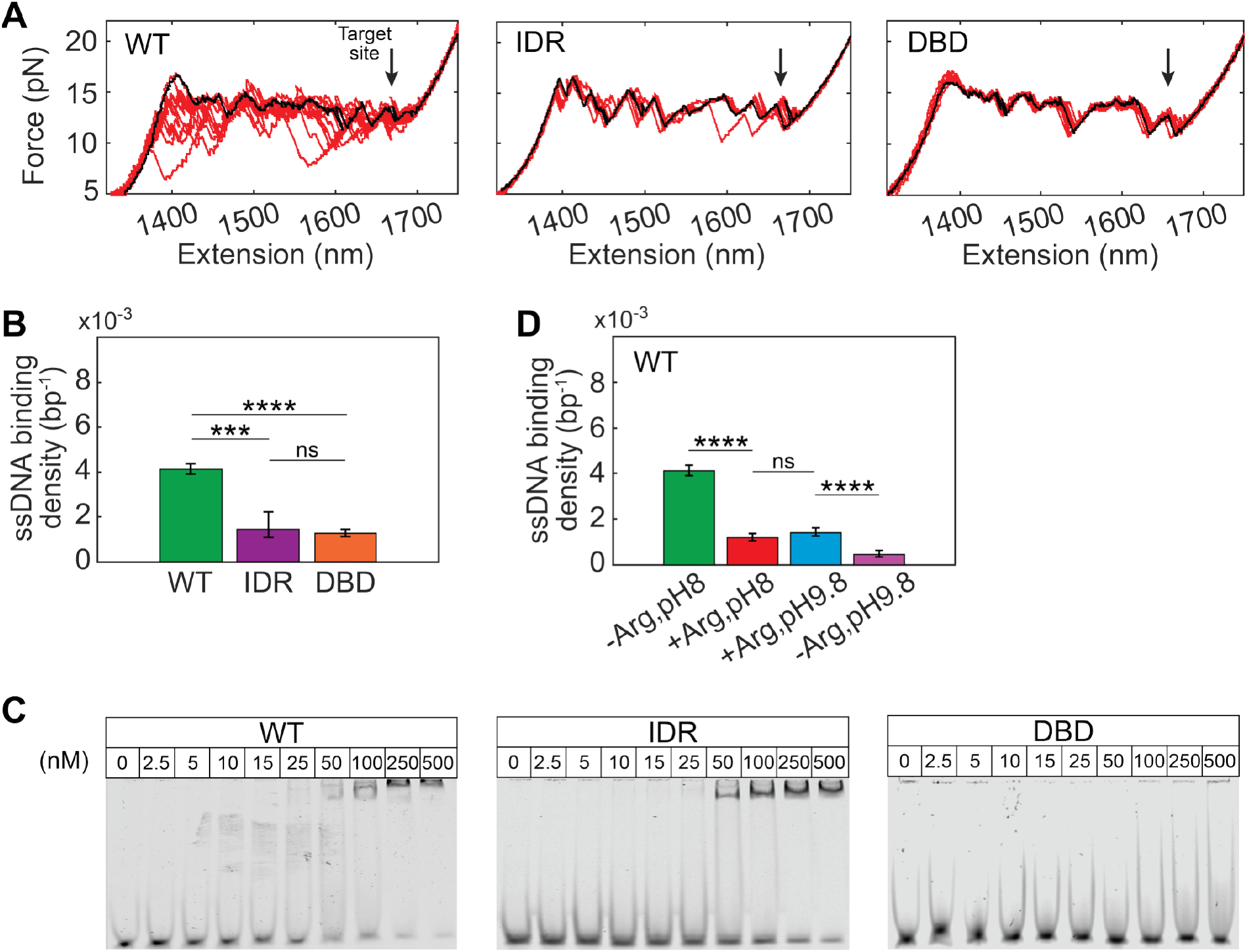
IDRs mediate Msn2 binding to single-stranded DNA. (A) Representative force-extension traces from DNA rezipping experiments with Msn2 variants. The black trace was obtained in a protein-free solution for reference. Traces with WT Msn2 show significant hysteresis due to non-specific binding to ssDNA at positions away from the binding site (marked by an arrow). (B) Quantification of ssDNA non-specific binding density, comparing Msn2 variants (total position bins n_WT_=8959, n_IDR_=806, n_DBD_=8773). Data shown as mean ± SEM, ***P<0.001, ****P<0.0001, χ^2^ test. (C) EMSA of fluorescently labeled ssDNA oligonucleotides with increasing concentrations of Msn2 variants. Bound complexes were observed for all variants except the DBD construct. (D) ssDNA non-specific binding density for the WT variant under different electrostatic perturbation conditions as described in Figure 1 (total position bins n_-Arg,pH8_=8959, n_+Arg,pH8_=8029, n_+Arg,pH9.8_=7068, n_-Arg,pH9.8_=8928). Data shown as mean ± SEM. ****P<0.0001, χ^2^ test. See also Table S8.

A surprising result from the kinetic experiments provided additional support for these results. Under conditions where DNA undergoes thermal breathing (Figure S5C), the distribution of open-state lifetimes is typically well described by a single exponent (Figure S5D, left). However, in the presence of WT, but not the DBD variant, a second population of long-lived open states emerged (Figure S5D, center and right), indicating transient stabilization of the open state by protein binding to ssDNA. Together, these results reveal a previously uncharacterized ability of Msn2 to interact with ssDNA through its disordered regions.

### IDRs enable sequence-dependent target search via 1D diffusion on DNA

Our findings that IDRs promote both non-specific DNA binding and faster association to specific motifs suggest a potential search mechanism: Msn2 may initially bind DNA non-specifically via its IDRs, then locate its target site by one-dimensional diffusion along the DNA. To directly test this model, we developed a new single-molecule assay, Sliding to Target Occupation (STO), designed to isolate this diffusion-based search mechanism (Figure 5A). In this assay, the DNA construct is fully unzipped, effectively eliminating the binding motif, and incubated in a TF-containing solution for one minute. Using our laminar flow cell, the construct is then transferred into a TF-free channel, where no additional proteins can bind the DNA and dissociation is irreversible. The construct then undergoes repeated unzipping and rezipping cycles to monitor for specific site occupation. Remarkably, following incubation with WT Msn2, we observed specific binding events at the motif (Figure 5B and Figure S6A, top) after a variable number of post-incubation cycles. No binding events were detected without prior incubation, indicating that Msn2 reaches the binding site by initially binding non-specifically elsewhere on the DNA and remaining associated through multiple unzipping and rezipping cycles while performing DNA-mediated diffusion. When IDRs were compromised, either by truncation or electrostatic screening, specific binding events following incubation decreased sharply (Figure 5C, Figure S6B). These results provide direct evidence for an IDR-facilitated search mechanism that depends on charge-mediated interactions.

**Figure 5:**
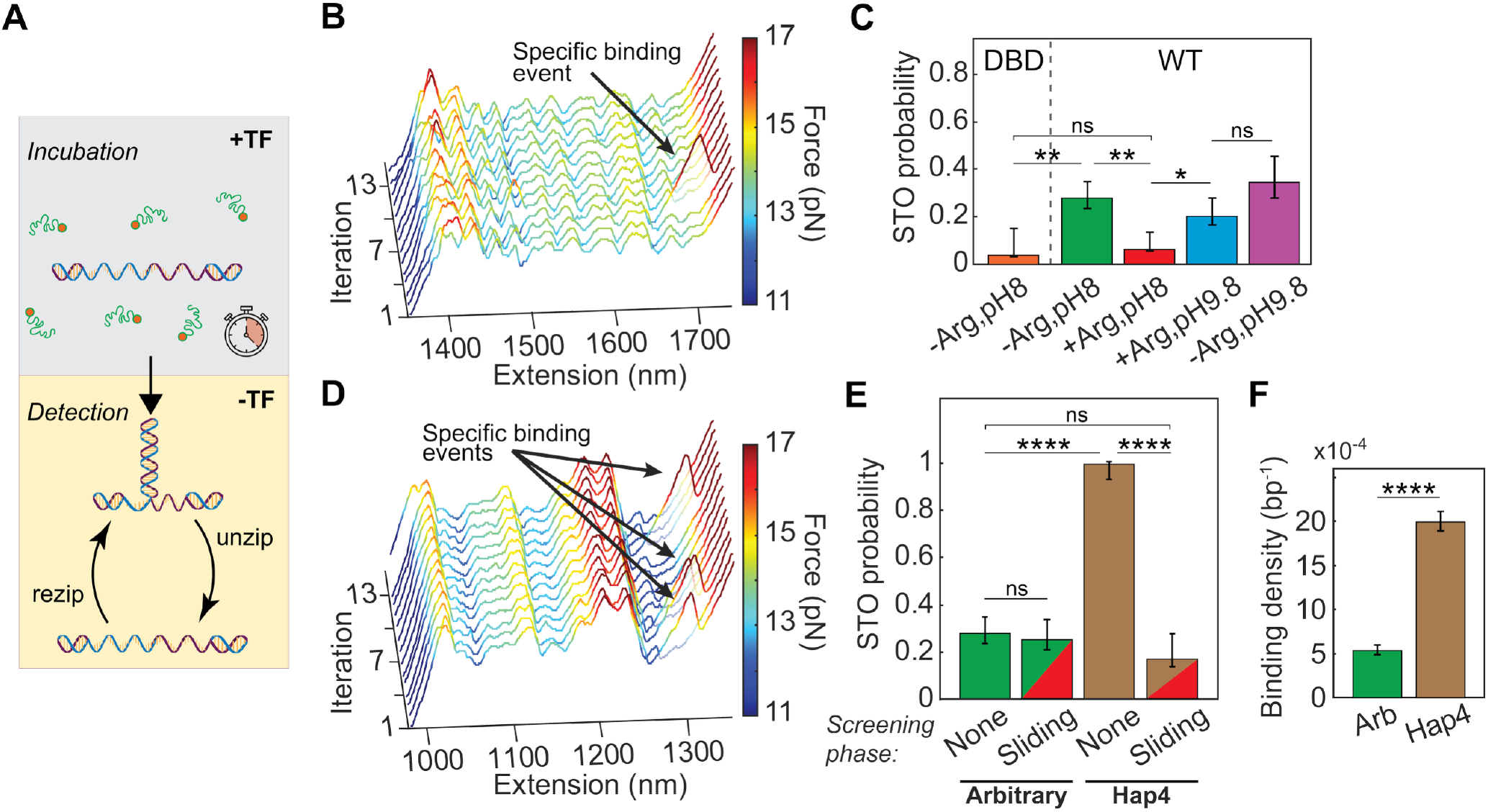
IDRs enable sequence-dependent target search via 1D diffusion on DNA. A) Schematic of the Sliding-to-Target Occupation (STO) assay. During the incubation phase, fully unzipped DNA is exposed to transcription factors, allowing non-specific loading. The construct is then transferred to a protein-free channel for the detection phase, where repeated unzipping– rezipping cycles monitor whether proteins reach the binding site via 1D diffusion. (B) Representative unzipping force-extension traces from the detection phase following a one-minute non-specific incubation with WT Msn2. A specific binding event at the target motif (arrow) appears during the fourth iteration. Each cycle lasts ∼8 seconds. (C) STO probability, defined as the fraction of DNA constructs showing specific binding events following non-specific incubation. IDR functionality was assessed by comparing WT Msn2 to either the DBD variant or the WT protein under electrostatic perturbation conditions as described in Figure 1 (total DNA molecules probed n_DBD_=27, n_-Arg,pH8_=61, n_+Arg,pH8_=49, n_+Arg,pH9.8_=50, n_-Arg,pH9.8_=26). Data shown as mean ± SEM, *P<0.05, **P<0.01, χ^2^ test. (D) Representative force-extension traces during the detection phase using a DNA construct in which the flanking sequence was replaced with a segment from the Hap4 promoter. Specific binding events occurred more frequently than in the arbitrary-sequence environment. (E) WT Msn2 STO probability comparison between different sequence environments flanking a constant binding site, with (split color bar) and without (solid color) electrostatic screening (50 mM L-arginine) applied during the sliding phase only. Left: arbitrary-sequence environment (green bar is same as -Arg,pH8 condition in C), (total DNA molecules probed n_None_=61, n_Sliding_=51). Right: Hap4 promoter (total DNA molecules probed n_None_=50, n_Sliding_=30). Data shown as mean ± SEM, ****P<0.0001, χ^2^ test. (F) Non-specific binding density for WT Msn2 with arbitrary sequence and Hap4 promoter sequence (total position bins n_Arbitrary_=40486, n_HAP4_=21390). Data shown as mean ± SEM, ****P<0.0001, χ^2^ test. See also Table S9.

To rule out the possibility that STO binding events arise solely from extended binding to ssDNA, we repeated the experiments performing the incubation step in a closed dsDNA state (i.e. without unzipping prior to incubation). This yielded similar STO probabilities (Figure S6C), indicating that neither the presence of the motif nor the formation of extended ssDNA during incubation is necessary for successful site localization during the diffusion phase. Notably, following ssDNA incubation with WT, we observed numerous binding events during the first rezipping iteration, but most vanished in subsequent cycles (Figure S6A, bottom). This pattern suggests that while a substantial population of Msn2 molecules initially binds to ssDNA during incubation, only a small subpopulation, potentially stabilized by additional DBD–DNA contacts, remains bound across multiple cycles and ultimately reaches the target motif.

We next asked whether this IDR-mediated search mechanism might contribute to Msn2’s ability to selectively occupy its motif only in certain promoters *in vivo*^25^. We replaced our standard (‘arbitrary’) DNA environment with a motif-free segment from the *Hap4* promoter, which is selectively bound by Msn2 *in vivo*. Importantly, the motif-containing segment was kept constant to avoid proximal flanking sequence effects. WT Msn2 showed a markedly higher probability of STO binding on the *Hap4* sequence compared to the arbitrary one (Figure 5D–E, S7A), and the specifically bound TFs were typically detected in earlier cycles (i.e. shorter times) (Figure S7B), indicating that features of the promoter environment accelerated target site localization.

We also observed that the *Hap4* promoter sequence induced significantly more non-specific binding than the arbitrary sequence, as reflected both in the density of non-specific events (Figure 5F) and in the fraction of molecules showing such events (Figure S7C). These non-specific interactions were also mechanically stronger, as reflected in their higher rupture forces (Figure S7A). When tracking these events over successive unzipping– rezipping cycles, we found their density remained approximately constant for both sequences (Figure S7D). Since the sliding phase of the STO assay occurs in a protein-free channel, these persistent interactions cannot result from non-specific re-binding events from the solution. Thus, the majority of Msn2 molecules that initially bind non-specifically to DNA, and survive the first rezipping cycle, persist on the DNA between consecutive unzipping and rezipping cycles. This suggests that the initial non-specific binding, mediated by the IDRs and stabilized by the DBD, is sequence-sensitive, but the dissociation rate is not. Supporting this interpretation, the density of non-specific binding events is non-uniform along the DNA sequence (Figure S7E). The sequence features that promote non-specific Msn2 interactions remain to be determined.

Finally, to test whether the diffusion step is sequence dependent too, we added L-arginine selectively to the TF-free channel, aiming to perturb IDR interactions during the diffusion phase without affecting initial non-specific binding. Under these conditions, both STO probability and detection times on the arbitrary sequence remained unchanged (Figure 5E, Figure S7B). In contrast, the *Hap4* promoter exhibited a sharp reduction in STO probability and a shift toward longer detection times, reaching both probability and detection time values comparable to those observed for the arbitrary sequence (Figure 5E, Figure S7B). These results suggest that while sliding can occur at a basal level with compromised IDRs activity, its sequence-specific enhancement requires IDRs–DNA interactions. Together, these findings highlight a role for IDRs in enabling promoter-specific search by contributing to both initial DNA engagement and preferential exploration of favorable sequences during one-dimensional diffusion.

## DISCUSSION

While canonical DBDs recognize short core sequence motifs, these motifs are typically degenerate and widely distributed throughout the genome. Yet *in vivo*, TFs occupy only a small fraction of these potential sites, suggesting that additional mechanisms confer functional specificity. This raises a fundamental question: how do TFs selectively bind their functional targets within a dense background of decoys? Motivated by *in vivo* studies showing that IDRs contribute to promoter specificity^4,25^, we used single-molecule optical tweezers to investigate the molecular mechanisms underlying this selectivity, using the yeast TF Msn2 as a model. Our findings reveal a multifaceted role for IDRs in supporting efficient and selective DNA search, demonstrating that direct interactions between disordered regions and DNA play a central role in conferring sequence specificity through mechanisms beyond canonical motif recognition.

Our results support a model in which Msn2 engages with DNA through a stepwise process facilitated by its disordered regions (Figure 6), which ultimately increases the association rate to the cognate motif by providing an additional binding pathway separate from direct binding from the solution. First, the IDRs mediate initial binding, allowing Msn2 to associate with DNA in the absence of a canonical motif. Once bound, Msn2 scans the DNA via one-dimensional diffusion, ultimately leading to recognition and stable binding at the canonical AGGGG motif. IDRs enhance this diffusion, thereby promoting more efficient and selective occupation of target sites. A central insight from our study is that both initial binding and diffusion are sequence sensitive. DNA derived from the *Hap4* promoter, which is selectively bound by Msn2 *in vivo*, enhances the formation of non-specific complexes and increases the probability and speed of site localization. These effects are abolished when the IDRs are removed. Site localization is also affected when the diffusion phase is perturbed without affecting initial binding, demonstrating that features of the promoter environment modulate both the likelihood of initiating a productive search trajectory and the efficiency with which the target is reached. Thus, sequence-selectivity emerges not only at the recognition site itself, but throughout the entire search process, and is encoded in the dynamic interactions between Msn2’s domains and the surrounding DNA. Notably, the observed sequence-selectivity, induced by replacing ∼300 bp of flanking sequence while keeping the binding motif constant, indicates that TFs’ search efficiency can be modulated over a short, ∼100 nm range. This spatial scale is below the resolution of fluorescence-based single-particle tracking approaches, underscoring the unique ability of single-molecule mechanical assays to resolve localized sequence effects on transcription factor dynamics. This model aligns with the classic facilitated diffusion framework for transcription factor target search^11,28,29^, and observations of 1D diffusion for various transcription factors^30,31^, while introducing an additional layer by demonstrating the critical role of disordered regions in this process in the case of Msn2.

**Figure 6:**
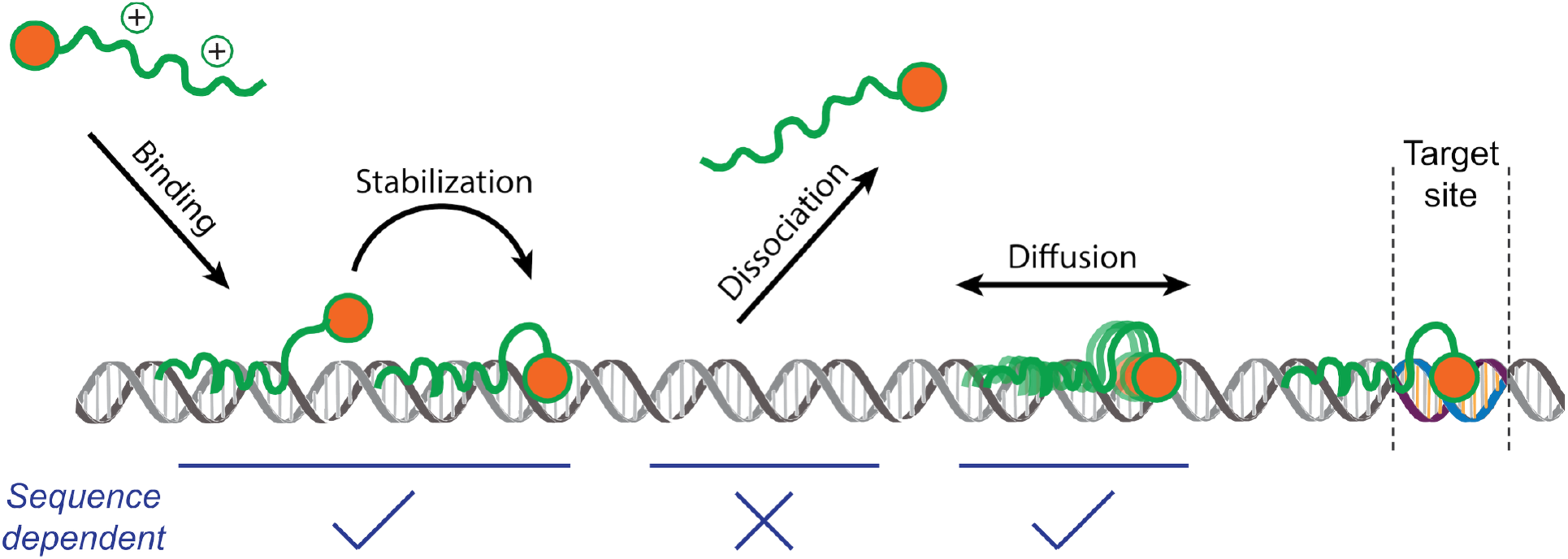
An emerging model of the IDRs-mediated searching mechanism of Msn2. Schematic representation of how Msn2 uses its intrinsically disordered regions (IDRs) to facilitate DNA binding and selective target localization. The process begins with (1) initial cooperative non-specific binding, driven by electrostatic interactions between the positively charged IDRs and the DNA, followed by (2) stabilization via the DNA-binding domain (DBD). Binding, stabilization, or both contribute sequence selectivity. Next, (3) one-dimensional diffusion along the DNA, modulated by the IDRs in a sequence-sensitive manner. Finally, (4) the protein stably binds the specific recognition motif. This mechanism illustrates how IDRs tune both the kinetics and spatial specificity of transcription factor target search, thereby contributing to promoter selectivity.

We show that disordered and structured regions cooperate to enable stable DNA engagement away from the canonical binding motif. While the IDRs mediate non-specific interactions with DNA, these alone were insufficient to generate detectable unzipping signatures in our single-molecule assay. In contrast, full-length Msn2, which contains both IDRs and the DBD, was detected at non-motif sites. This suggests that the DBD, even outside its cognate sequence, provides additional stabilization that cooperates with the IDRs to form strand-engaging complexes. Rather than acting independently, the disordered and structured domains function as an integrated unit: the IDRs promote charge-mediated initial contact with the DNA, while the DBD reinforces these interactions, likely through weak hydrogen bonding or van der Waals contacts. Notably, while we refer to the initial binding as “non-specific,” it exhibits clear sequence preference, as it was significantly increased for the *Hap4* promoter. This increase may arise from differential association by the IDRs, selective stabilization by the DBD, or both. In the case of the IDRs, several biophysical mechanisms could support such selectivity. First, DNA sequences generate distinct electrostatic landscapes, with local charge variations that may bias initial IDR contact^32,33^. Second, sequence-dependent mechanical properties could affect how IDRs conform around the DNA backbone^34^. Third, certain DNA regions may exhibit enhanced base-pair breathing or transient melting, increasing access to exposed bases. This last possibility is particularly intriguing in light of our finding that Msn2’s IDRs can bind to single-stranded DNA.

Perturbation experiments helped clarify the molecular interactions through which IDRs contribute to the search process. The addition of 50 mM free L-arginine to the buffer, positively charged at physiological pH, led to a marked reduction in both non-specific binding and specific site engagement, in equilibrium and STO assays alike. These effects are consistent with charge-based screening of attractive IDR–DNA interactions. Although L-arginine could in principle affect binding through other mechanisms, these are unlikely to account for the magnitude of the observed effects at the concentrations used. Moreover, control experiments showed that increasing ionic strength with KCl or raising the pH to 9.8, which neutralizes the amino group of L-arginine, reversed the inhibitory effect. These findings highlight the critical role of charge-mediated interactions in enabling the IDR-mediated search mechanism, particularly by facilitating initial binding and DNA association. However, they do not imply that electrostatics underlie the *sequence-dependent* aspects of the search, whose molecular basis may involve additional interactions, such as hydrophobic contacts, previously shown *in vivo* to contribute promoter-specific occupancy^1^.

Our observation that Msn2 can bind to ssDNA through its IDRs, supported by the hysteresis observed during rezipping experiments and the extended open periods in thermal fluctuation measurements, represents a novel finding with potential functional implications. Transcriptionally active regions often contain transient ssDNA^35^. The capacity of IDRs to engage with these structures could potentially contribute to transcription factor recruitment to active genes or influence the dynamics of transcriptional regulation. Recent work^36^ demonstrated that the intrinsically disordered C-terminal domain of linker histone H1 exhibits preferential binding to single-stranded nucleic acids and can undergo liquid-liquid phase separation upon binding. This suggests that IDR-mediated interactions with single-stranded nucleic acids may represent a common functional modality across diverse DNA-binding proteins. Notably, the flexible and dynamic nature of ssDNA may promote non-electrostatic contacts, such as hydrogen bonding and potentially π–π or cation– π interactions involving aromatic or basic residues. This is supported by structural analyses of RNA–protein complexes^37^, suggesting that similar interactions could contribute to IDR engagement with single-stranded nucleic acids.

While our current study focuses on DNA binding in a purified system, transcription factors must navigate a considerably more complex chromatin environment *in vivo*. Nucleosomes represent a major barrier to TF binding, potentially occluding binding sites and impeding the one-dimensional diffusion process characterized here. Interestingly, *in vivo* studies have shown that IDRs direct Msn2 to promoters with “fuzzy” nucleosome architectures^4^, suggesting a possible link between disordered regions and dynamic chromatin states. Our previous work showed that nucleosomes modulate the energy landscape for TF binding through multiple mechanisms, including site accessibility and nucleosome dynamics^38–40^. Others have shown that the presence of nucleosomes can alter TF-DNA interactions, modulating TF-DNA recognition beyond simple steric occlusion^41^. Furthermore, the electrostatic and potentially hydrophobic interactions mediated by IDRs may enable Msn2 to associate with the histone octamer surface or acidic histone tails. The IDR-mediated search mechanism we describe could thus be particularly relevant in this chromatin context, as it could facilitate the scanning of partially accessible DNA regions during transient nucleosome unwrapping events, or facilitate engagement with both DNA and chromatin components during the search process.

Finally, given the prevalence of intrinsic disorder in eukaryotic transcription factors_42_, the search mechanism identified here may represent a fundamental principle that has been repeatedly exploited throughout evolution. While DBDs typically change through discrete mutational events that alter specific base contacts, IDRs can evolve more rapidly due to their tolerance for a wider range of sequence variations, potentially accelerating the evolution of gene regulatory networks^43^. Our findings suggest that modifications to IDRs could tune not only the activation potential of TFs (as previously established) but also their DNA search properties and target specificity. If so, the disordered architecture of transcription factors may be as critical to their regulatory specificity as their structured DNA-binding domains, challenging conventional views of sequence-specific DNA recognition.

### Limitations of the study

While our study provides mechanistic insight into the role of IDRs in Msn2-mediated DNA binding and target search, several limitations should be considered. First, our experiments were conducted *in vitro* using purified components, which do not fully recapitulate the complexity of the nuclear environment *in vivo*. Chromatin structure, nucleosome organization, and interactions with other DNA-binding proteins or cofactors likely influence Msn2 binding dynamics in ways not captured by our experimental system. Second, while we observed clear sequence dependence in the IDR-mediated search process, we examined only a small set of DNA sequences. A more comprehensive analysis would be required to identify broader sequence determinants or develop predictive models of IDR-mediated targeting across the genome. Finally, although we propose that these findings may extend to other IDR-containing transcription factors, we tested only Msn2. Given the substantial diversity in IDR sequence composition, charge distribution, and structural plasticity across TFs, it remains unclear to what extent the specific mechanisms identified here are generalizable.

## RESOURCE AVAILABILITY

### Lead contact

Further information and requests for resources and reagents should be directed to and will be fulfilled by the lead contact, Ariel Kaplan.

### Materials availability

All unique reagents generated in this study are available by request from the lead contact.

### Data and code availability

Any additional information required to reanalyze the data reported in this paper is available from the lead contact upon request.

## ACKNOWLEDGMENTS

We thank Hagen Hofmann, Naama Barkai and Arnon Henn for valuable comments, and Gabriel Rosenblum and Vladimir Mindel for their assistance.

This work was supported by the Israel Science Foundation (Grant 937/20 to AK).

## AUTHOR CONTRIBUTIONS

A.K. and N.S. conceived the project. N.S., C.B., and M.G. designed and performed all experiments. N.S., C.B., and H.K. analyzed the data. N.S., C.B., S.H. and N.N. prepared experimental materials. N.S. and C.B. prepared the figures. A.K. and N.S. wrote the manuscript. A.K. supervised the project.

## METHODS

### Expression and Purification of Msn2 Proteins

All three Msn2 proteins were expressed using pET-24(b+) vector in *E. coli* BL21 (DE3) cells. The details of the constructs are presented in Table S1. The WT and IDR (N-terminal His6-tagged) were purified from inclusion bodies (IBs), whereas the DBD (N-terminal His6-MBP-tagged) was purified from the soluble fraction.

For the WT and IDR, the bacteria were grown in 4 l LB medium containing 50 µg/ml kanamycin, 0.5% glucose, and 4 g/l serine at 37 °C (250 rpm). The cultures were induced with 1.0 mM IPTG at OD_600_ 0.8 and incubated overnight under the same conditions. The cells were harvested by centrifugation (3,000 × *g*, 30 min, 4 °C), and the cell pellets were suspended in ice-cold lysis buffer (50 mM Tris-HCl pH 8.0, 10 mM EDTA, Roche cOmplete protease inhibitor cocktail tablet (1 for each 15 ml). After sonication (70% amplitude, pulse on 2 s, pulse off 10 s, 60 cycles, on ice) and centrifugation (48,200 × *g*, 30 min at 4 °C), the precipitate was mixed with 60 mM EDTA, 6% Triton and 1.5 M NaCl, homogenized, and incubated for 30 min at 4 °C to solubilize membrane debris. The resulting suspension was centrifuged (48,200 × *g*, 30 min at 4 °C) and the IB precipitate was re-suspended in 100 mM Tris-HCl pH 8.0 and 1 mM EDTA, homogenized, and centrifuged using the same conditions. This IB wash was repeated 2 times. Finally, the IB pellet was re-solubilized in 50 mM Tris-HCl pH 7.5, 10 mM imidazole, 6 M GdmCl, 100 mM DTT and 10 mM EDTA for more than 2 h at 4 °C. The supernatant was collected by centrifugation (48,200 × *g*, 30 min at 4 °C) and the buffer was exchanged to 50 mM Tris-HCl pH 7.5, 6 M GdmCl, and 10 mM imidazole, using a HiPrep desalting column (26/10, GE) to remove DTT and EDTA. The protein-containing fraction was then loaded on a HisTrap Ni-NTA column (5 ml, Cytiva) and the protein was eluted with 50 mM Tris-HCl pH 7.5, 6 M GdmCl, and 0.5 M imidazole. To remove the His6-tag, the eluted protein was supplemented with HRV3C protease (0.15 mg/ml) and the solution was first dialyzed against 125 volumes of cleavage buffer (50 mM Tris-HCl, 300 mM KCl, 300 mM L-Arg and 1 mM DTT, measured pH 7.5) for 1 h (buffer was changed after 30 min) at 25 °C and then against 200 volumes for overnight at 4 °C. GdmCl and imidazole powders (to final concentration of 6 M and 10 mM, respectively) were added to the cleavage reaction and loaded on the HisTrap column. Cleaved protein was obtained in the flow-through, while the His6-tag and HRV3C protease were retained on the column. The cleaved protein was then reduced with 30 mM DTT, and further purified either by size-exclusion chromatography (SEC, Superdex 200 Increase 10/300 GL, Cytiva) and the protein-containing fractions were exchanged to storage buffer (20 mM Tris-HCl pH 8 and 150 mM KCl) using a HiTrap desalting column (5 ml, Cytvia), or passed it through a reverse-phase HPLC Jupiter C4 column and then dissolved in the storage buffer. Purity of the proteins was confirmed by SDS-PAGE (Figure S1). The protein concentration was determined using a predicted extinction coefficient of *ε*_280_ = 24,870 M^−1^ cm^−1^ (both for WT and IDR) and stored at −80 °C.

For the DBD, bacteria were grown in 4 l LB medium containing 50 µg/ml kanamycin and 0.5% glucose at 37 °C (250 rpm). The cultures were induced at OD_600_ 0.8 with 0.5 mM IPTG and incubated at 25 °C (250 rpm) for an additional 8 h. The cells were harvested by centrifugation (3,000 × *g*, 30 minutes, 4 °C), and cell pellets were suspended in ice-cold lysis buffer (50 mM Tris-HCl pH 8, 10 mM EDTA, Roche cOmplete protease inhibitor cocktail tablet (1 for each 15 ml). After sonication (70% amplitude, pulse on 2 s, pulse off 10 s, 60 cycles, on ice) and centrifugation (48,200 × *g*, 30 min at 4 °C), the supernatant was collected and reduced with 50 mM DTT on ice for 1 h. In the next step, the supernatant was loaded on an amylose column (resin from NEB, packed in GE XK 16/40 column), washed with 20 mM Tris-HCl pH 7.5, 50 mM NaCl, 1 mM CaCl2 and the desired protein was eluted with 20 mM Tris-HCl pH 7.5, 50 mM NaCl, 1 mM CaCl2 and 10 mM maltose. To remove the DNA impurities, the protein was first reduced with 50 mM DTT on ice for 1 h, then diluted 5-fold with 15 mM Tris-HCl pH 7.3, 2 mM EDTA, and loaded on a DEAE FF ion-exchange column (5 ml, Cytiva). The flow-through (containing most of the protein) was collected, concentrated and then the buffer was exchanged to 20 mM Tris-HCl pH 7.5, 10 mM imidazole and 1.5 M NaCl, using the HiPrep desalting column. Afterwards, the solution was loaded on a 5 ml HisTrap column, washed with 10 column volume (CV) of 20 mM Tris-HCl pH 7.5, 25 mM imidazole and 0.5 M GdmCl, re-equilibrate with 20 mM Tris-HCl pH 7.5, 10 mM imidazole and 1.5 M NaCl, and finally eluted with 20 mM Tris-HCl pH 7.5, 0.5 M imidazole, and 1.5 M NaCl using a linear a gradient from 0 to 100% over 5 min (3 ml/min). The protein fractions were pooled and the buffer was exchanged to 20 mM Tris-HCl pH 7.5 and 100 mM NaCl using the HiTrap desalting column. To remove the MBP-His6-tag, the protein solution was incubated with 0.15 mg/ml HRV3C protease for 2 h at 25° C and passed it through the reverse-phase HPLC Jupiter C4 column. The pure lyophilized DBD (as confirmed by SDS-PAGE, Figure S1) was dissolved in storage buffer (20 mM Tris-HCl pH 7.5 and 100 mM NaCl), the concentration was measured using bicinchoninic acid (BCA) assay (because DBD has a computed extinction coefficient of *ε*_280_ = 0 M^−1^ cm^−1^) and stored at −80 °C.

### Single-molecule experiments

The constructs for single-molecule experiments were generated as previously described^27,40^ with some changes. All constructs were assembled by ligation of several DNA segments (Figure S8): A pair of DNA “handles”, an “upstream environment” segment (UE, ∼300 bp), a “binding region” segment (BR), and a short “hairpin” segment (HP). For the kinetics experiments in Figure 2, since the DNA is unzipped up to the binding site itself, an additional “downstream environment” segment (DE, ∼250 bp) was inserted between the BR and HP, to create a downstream flanking sequence similar to the upstream one. The two ∼2000 bp DNA “handles” were prepared as previously reported^27,39^, each incorporating a specific tag (biotin or digoxygenin) on one end and a complementary ssDNA overhang on the other, and were then annealed. The UE was amplified from the *Cga* gene promoter using mouse genomic DNA as a template or, for the experiments in Figure 5, from the *Hap4* gene promoter using *S. Cerevisiae* genomic DNA as a template. Primers are listed in Table S15. PCR products were digested with DraIII-HF (R3510L; NEB) according to the manufacturer’s instructions, and purified using a QIAquick PCR Purification Kit (28106; QIAGEN). BR for the experiments described in Figure S2A is a ∼600 bp segment from the *Hor7* gene promoter that includes four Msn2 binding motifs, and was produced via PCR with the pET24b-Hor7 plasmid (kindly provided by Dr. H. Hofmann) as a template (primers listed in Table S15). The product was digested with SFI-I (R0123S; NEB). In all other experiments, BR is a 60 bp AT-rich sequence flanking a single Msn2 binding motif and was produced by annealing two ssDNA oligonucleotides (Table S16), designed with overhangs at their ends in preparation for their ligation assembly. Complementary ssDNA oligos were first phosphorylated using T4 Polynucleotide Kinase (M0201L; NEB), according to the manufacturer’s instructions, mixed in a 1:1 molar ratio in T4 DNA Ligase Reaction Buffer (B0202S; NEB), incubated at 90°C for 5 minutes, and cooled down slowly to room temperature. BR for the Egr1-based experiment, shown in Figure S3B, is a 68 bp sequence designed with a single Egr1 binding motif along with a 5 bp native flanking segment from the *Lhb* gene promoter, and was produced in a similar annealing method. DE was amplified from the *Cga* gene promoter by PCR with mouse genomic DNA (primers listed in Table S15) and digested with DraIII-HF (R3510L; NEB). HP is a 27 nt ssDNA molecule that folds into a hairpin with a 10 bp stem (Table S16), and contains a 3 nt overhang to facilitate ligation (TGC for the kinetic experiments, CTA for all other experiments).

To assemble the full construct, the annealed handles were ligated to UE in a 1:5 reaction overnight with T4 DNA ligase (M0202L; NEB), and the ligation product was gel-purified. Separately, BR was ligated to HP in a 1:5 reaction overnight with T4 DNA ligase (M0202L; NEB). Handles-UE and BR-HP were then ligated in a 1:5 reaction for 30 minutes at room temperature, using Rapid ligase (C671B; Promega). For the kinetic experiments (Figure 2), DE was ligated to HP in a 1:5 reaction overnight with T4 DNA ligase (M0202L; NEB), and then a three-piece ligation with Handles-UE, BR and DE-HP, in a 1:5:25 ratio, was performed for 30 minutes at room temperature, using Rapid ligase (C671B; Promega).

The full construct was incubated for 15 minutes on ice with 0.8 μm polystyrene beads (Spherotech), coated with anti-Digoxigenin (anti-Dig) molecules. The reaction was then diluted 1000-fold in binding buffer (BF): 20 mM Tris·HCl pH 8 (or 9.8, as specified), 5 mM MgCl_2_, 1 mM TCEP, 3% v/v glycerol and 0.01% BSA, 0.001% Tween20, 1uM EDTA, 2uM ZnCl, 150 mM KCl (unless otherwise indicated), and (when indicated) 50mM L-arginine. The binding buffer for Egr1 experiments was composed of 10 mM Tris·HCl pH 7.4, 150 mM NaCl, 1.5 mM MgCl_2_, 1 mM DTT, 3% v/v glycerol, and 0.01% BSA. Tether formation was performed *in situ* (inside the optical tweezers’ experimental chamber) by trapping a DNA-bound anti-Dig bead in one trap, 0.9 μm streptavidin-coated polystyrene beads in the other trap, and bringing the two beads into proximity to allow binding of the biotin tag to the streptavidin-coated bead.

All the experiments with the Msn2 variants were conducted at 50 nM protein concentration, except for the kinetics experiments, which were conducted at 10 nM. Experiments with Egr1 were conducted at 4 nM.

### Optical Tweezers

Experiments were performed in a custom-made dual-trap optical tweezers apparatus, as previously reported^26,27,44^. Briefly, the beam from an 852 nm laser (TA PRO, Toptica) is coupled into a polarization-maintaining single-mode optical fiber. The collimated beam out of the fiber is split by a polarizing beam splitter (PBS) into two orthogonal polarizations, each directed into a mirror and combined again with a second PBS. One of the mirrors is mounted on a nanometer-scale mirror mount (Nano-MTA, Mad City Labs). An X2 telescope expands the beam while imaging the plane of the mirrors into the back focal plane of the focusing microscope objective (Nikon, Plan Apo VC 60X, NA/1.2). Two optical traps are formed at the objective’s focal plane, each by a different polarization, and with a typical stiffness of 0.3-0.5 pN/nm. The light is collected by a second, identical objective, and the two polarizations are imaged onto two Position Sensitive Detectors (First Sensor), after being separated by a PBS. The instantaneous position of the beads relative to the center of their traps is determined by back focal plane interferometry^45^. Calibration of the setup is done by analysis of the thermal fluctuations of the trapped beads^46^, which are sampled at 100 kHz. Experiments were conducted using a 5-channel laminar flow cell (Lumicks), which was passivated following a published protocol47 with some modifications. Briefly, we washed the chamber twice by flushing alternately 1M NaOH and 1% Liquinox for 10 min each. Casein (1%) was sonicated, filtered, diluted to 0.2%, and flushed into the chamber, where it was incubated for 40 min. After the incubation, the system was washed with binding buffer.

### Data analysis

The fundamental analysis of the acquired data followed the same methodology previously described^27,40^. Briefly, data signals were acquired at 2500 Hz and stored for analysis using Matlab (MathWorks) scripts. The data was converted into force and extension vectors using the experimentally determined calibration parameters. When indicated (Figures 1, S2) the position (or number of bp unzipped) was calculated by subtracting the stretching of the dsDNA and ssDNA parts of the construct, which we calculate using extensible-worm-like-chain (eWLC)^48^ and worm-like-chain (WLC) models, respectively.

### Motif-specific binding experiments

Motif-specific binding events were identified as events where the maximal force *Fi* at the nominal binding site position (±5 bp) exceeded a threshold force. This threshold was determined as the average maximal force at the binding site position from multiple traces taken in a protein-free solution, plus three times the standard deviation of these force values. When applying the threshold criteria to the data obtained from the protein-free solution, no binding events were detected. The binding probability was calculated as n/N, where n is the number of times a binding event was identified, and N is the total number of cycles. The uncertainty in the binding probability was calculated with the adjusted Wald interval of binomial proportions^49^, and the significance of differences was assessed using a χ^2^ test. The breaking force was calculated as the mean *Fi* for the cycles where binding was detected. The significance of the difference between bound complex breaking forces was evaluated with a two-tailed Student’s t-test. Differences were considered statistically significant when p < 0.05.

### Non-specific binding experiments

This experiment aims to detect non-specific binding events to either dsDNA or ssDNA, occurring over 20 bp away from the protein binding site motif. The maximal forces during unzipping iterations at a protein-free solution were measured throughout the DNA motif-flanking environment with 10 bp bins. This allowed us to set a threshold on the maximal force expected in the absence of a protein in a position-dependent manner, as three times the standard deviation above the averaged maximal force in each bin. The force data of each unzipping iteration, from experiments in the presence of the protein, was then compared, per bin, to the calculated threshold of that bin and marked as a dsDNA non-specific binding event if exceeded. Position-resolved non-specific binding density (Figure S7E) was calculated as the average number of bound events in each bin, divided by the bin size. The overall binding density was calculated by averaging over all the bins. Similarly, a minimal force threshold was calculated from the minimal force data of the rezipping iterations, to form a threshold on the minimal force expected in the absence of a protein, and identify ssDNA non-specific binding events during rezipping iterations. The ssDNA binding density was calculated in a similar manner to the dsDNA binding density. To avoid misidentified events at positions that are particularly noisy in terms of their maximal force, unzipping/rezipping traces taken in protein-free solution were also verified against the calculated thresholds per position bin. In case one of the tested DNA molecules showed above/below threshold behavior in a protein-free solution, the relevant position bins were excluded from the binding density calculation of the corresponding molecule measurements that were held in the presence of the protein. Finally, the “background” density calculated in a protein-free solution was subtracted. The significance of differences between non-specific binding densities was evaluated with a χ^2^ test. Differences were considered statistically significant when p < 0.05. In addition to this position-resolved method, an alternative method of quantifying non-specific binding was used in Figure S7C. The non-specific binding was calculated as the percentage of DNA molecules showing any non-specific binding event out of the total number of tested molecules. The significance of differences between these binding probabilities was evaluated with a χ^2^ test. Differences were considered statistically significant when p < 0.05.

### Sliding to target occupation (STO) experiments

The experiment aims to identify motif-specific binding events occurring in a protein-free solution following non-specific binding of the protein to the DNA construct. For that, separate channels (one protein-free channel and the second containing the variant under test) in our microfluidic laminar flow chamber (Lumicks) were utilized in a three-phase experiment. First, the DNA construct was fully unzipped in the protein-free channel. Next, it was translocated to the channel where the protein was present. At that stage, the unzipped construct had no effective binding site, hence proteins could only bind to it non-specifically. A 60-second incubation period was provided to allow stabilization of the non-specific binding process. Lastly, the construct was moved back to the protein-free chamber channel, where 20 unzipping-rezipping iterations were performed to probe for binding events. STO probability was calculated as the fraction of DNA constructs showing any specific binding event following non-specific incubation. The significance of differences between STO probabilities was evaluated with a χ^2^ test. Differences were considered statistically significant when p < 0.05.

### Kinetics experiments

Dissociation and association rates of the protein-DNA complex were calculated as previously described^27^, with some changes as detailed below. Briefly, fixing the positioning of the unzipping fork in the vicinity of the protein’s binding motif allows identifying protein binding by probing thermal fluctuations between “closed” (dsDNA) and “open” (two ssDNA) states, which are suppressed upon TF binding. After identifying closed and open stats using the HAMMY algorithm^50,51^, their durations were determined. Closed states longer than a predetermined threshold, chosen to minimize the overall misidentified states, were classified as “bound” states. Non-bound periods above 15 ms were defined as “unbound” states. Association and dissociation rates were calculated as the reciprocal of the average unbound and bound states duration, respectively. Statistical significance of differences in kinetic rates was assessed using a two-tailed non-parametric permutation test (10,000 iterations). In each iteration, dwell periods were randomly reassigned between the two condition groups to simulate the null hypothesis, and the rate difference was calculated. P-value was defined as the proportion of permuted rate differences with a magnitude greater than or equal to that of the observed difference. Dissociation rates for the two populations observed in Figure 2C were calculated by fitting the bound periods’ distribution with a double exponential function of the form *p* · *exp*(−*k*_*off*1_*t*) + (1 − *p*) · *exp* (−*k*_*off*2_*t*). Significance difference between the kinetic rates was assessed using a two-tailed non-parametric permutation test (10,000 iterations). Differences were considered statistically significant when p < 0.05.

### Electrophoretic mobility shift assays (EMSA)

A 51 bp dsDNA probe labeled with 5′ IRD800 was prepared by annealing 20 μM of each complementary oligonucleotide (IDT; sequences in Table S17) in STE buffer (100 mM NaCl, 10 mM Tris-HCl pH 8.0, 1 mM EDTA) in a final volume of 100 μL. The mixture was heated to 95°C for 5 min and then slowly cooled to room temperature overnight. Residual single-stranded DNA was removed by exonuclease treatment, and the resulting dsDNA was purified by ethanol precipitation. Binding reactions were performed in a 20 µl solution containing EMSA reaction buffer (20 mM Tris-HCl pH 8, 150mM KCl, 5 mM MgCl_2_, 1mM TCEP, 1µM EDTA, 2µM ZnCl_2_), 1 nM DNA and 2.5-500 nM of the purified Msn2 variants (WT, IDR, or DBD). Reactions were incubated at room temperature for 30 min and then resolved on a non-denaturing 6% polyacrylamide gel prepared and run in 0.5× TBE buffer (45 mM Tris-borate, 1 mM EDTA). Samples were loaded directly onto the gel, which was run at 100 V for 60 min at room temperature.

Gels were scanned using an Odyssey DLx imaging system (LI-COR), and band intensities were quantified using Image Studio Lite software. Signal intensities corresponding to the bound and unbound DNA fractions were extracted. Quantification was performed using a thermodynamic equilibrium model with two sequential binding steps. Under the assumption that the total DNA concentration is much lower than the total transcription factor (TF) concentration, the fractional occupancies were calculated as 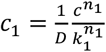 and 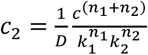, where 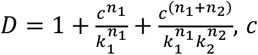 is the free TF concentration, *k*_1_ and *k*_2_ are the dissociation constants for the first and second binding steps, and *n*_1_ and *n*_2_ are their respective Hill coefficients. Model parameters were determined by non-linear least-squares fitting to the experimental data using MATLAB’s Curve Fitting Toolbox, with initial parameter estimates based on observed half-maximal binding concentrations. Goodness of fit was evaluated using the coefficient of determination (R^2^) and residual analysis.

### Mass-photometry

A 51 bp double-stranded DNA probe, identical to the one used in the EMSA experiments but lacking the fluorescent label, was prepared using the same annealing protocol (Table S17). Binding reactions were carried out in a 20 µl solution containing EMSA buffer, 50 nM WT Msn2, and 50 nM DNA, followed by 30 min of incubation at room temperature. Mass photometry measurements were acquired using a OneMP mass photometer (Refeyn Ltd, Oxford, UK). Microscope coverslips (No. 1.5, 24 × 50 cat 0107222, Marienfeld®) were cleaned by sequential sonication in 50% isopropanol (HPLC grade)/Milli-Q H2O, and Milli-Q H2O (5 min each), and dried using a clean nitrogen stream. Four-welled silicone gaskets (Re-useable culturewell™ gaskets 3mm diam x 1mm depth, cat GBL103250-10EA, Sigma-Aldrich®) were cut, washed with ethanol, followed by MilliQ water, and dried using a nitrogen stream. Dried gaskets were placed at the center of the coverslip, and each well was used for one measurement. PBS (ultra-pure, VWR®) buffer was first loaded into the well, and the focal position was identified and secured for the entire measurement with an autofocus system based on total internal reflection. Sample solutions were introduced into the well with the loaded buffer and mixed several times (final concentration 50nM) for each acquisition. Standard proteins Bovine Serum Albumin (mass: 66 kDa, 132 kDa; Thermo Scientific®) and Bovine Serum IgGs (mass: 150 kDa, 300 kDa; Sigma-Aldrich ®) were similarly measured and used to generate a linear molecular weight calibration curve. Following autofocus stabilization, movies of 120 seconds duration were recorded per sample. Data acquisition was performed using AcquireMP (Refeyn Ltd, 2024 R1.1) and data analysis using DiscoverMP (Refeyn Ltd, 2024 R1).

