## Supplementary Information for "Intrinsically disordered regions facilitate Msn2 target search to drive promoter selectivity"

**SUPPLEMENTARY FIGURES**

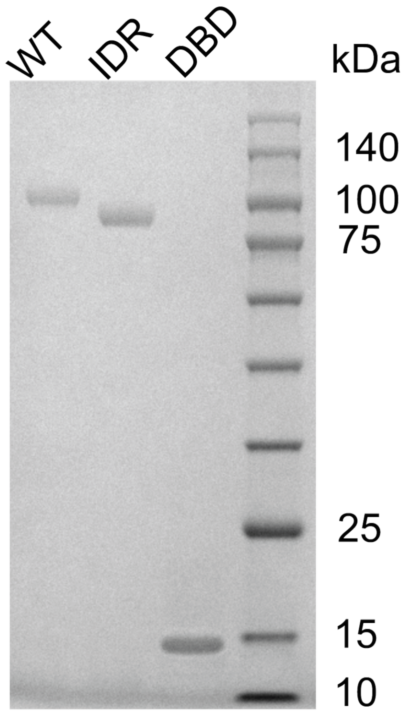


**Figure S1. Msn2 protein variants**. Coomassie-stained SDS-PAGE of the three protein variants following final purification. WT (78 kDa), IDR (71 kDa), and DBD (7.6 kDa), are shown relative to a protein molecular weight marker (PM2500, ExcelBand) on the right lane.

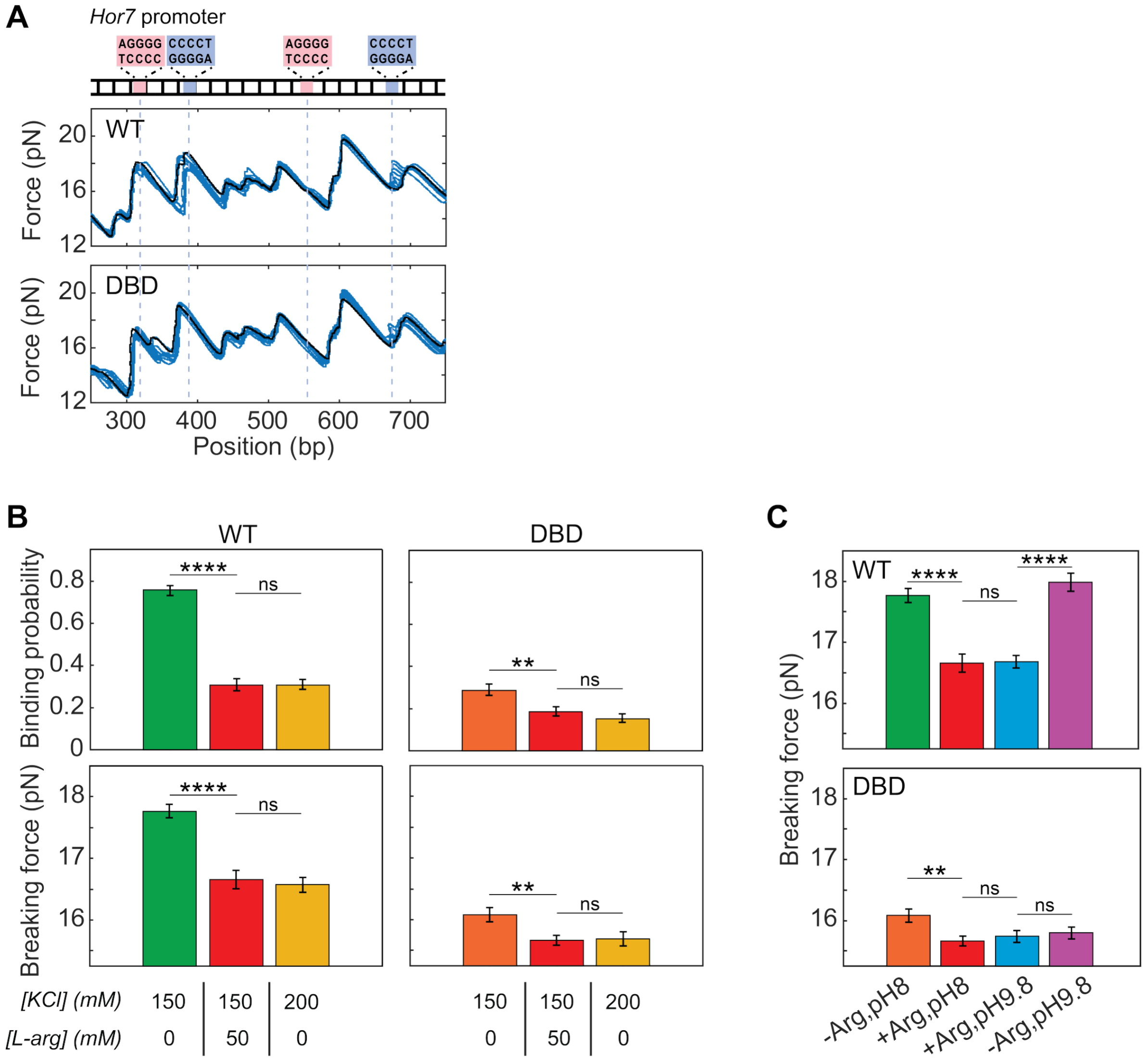


**Figure S2. Supporting results for IDRs contribution to the specific bound complex**. (A) Representative unzipping iterations in the presence of WT and DBD variants using the *Hor7* gene promoter, which contains four binding sites (marked with dashed lines). The black trace shows unzipping in a protein-free solution for reference. (B) Binding probability (total unzipping iterations $n_{WT;KCl150,Larg0}$=296, $n_{WT;KCl150,Larg50}$=258, $n_{WT;KCl200,Larg0}$=370, $n_{DBD;KCl150,Larg0}$=289, $n_{DBD;KCl150,Larg50}$=309, $n_{DBD;KCl200,Larg0}$=333) and breaking force ($n_{WT;KCl150,Larg0}$=226, $n_{WT;KCl150,Larg50}$=79, $n_{WT;KCl200,Larg0}$=114, $n_{DBD;KCl150,Larg0}$=83, $n_{DBD;KCl150,Larg50}$=57, $n_{DBD;KCl200,Larg0}$=50) for the WT and DBD variants, comparing two types of screening perturbation methods: adding 50mM free L-arginine and increasing [KCl] by 50mM. Data shown as mean ± SEM, **P<0.01, ****P<0.0001, $\chi^2$ test and Student's t-test, respectively. See also Table S10. (C) Breaking force for the AT-rich arbitrary-sequence environment, under different electrostatic perturbation conditions as described in Figure 1, for the WT (top panel; total unzipping iterations $n_{-Arg,pH8}$=226, $n_{+Arg,pH8}$=79, $n_{+Arg,pH9.8}$=155, $n_{-Arg,pH9.8}$=166) and DBD variants (bottom panel; $n_{-Arg,pH8}$=289, $n_{+Arg,pH8}$=83, $n_{+Arg,pH9.8}$=57, $n_{-Arg,pH9.8}$=73). Data shown as mean ± SEM, **P<0.01, ****P<0.0001, Student's t-test.

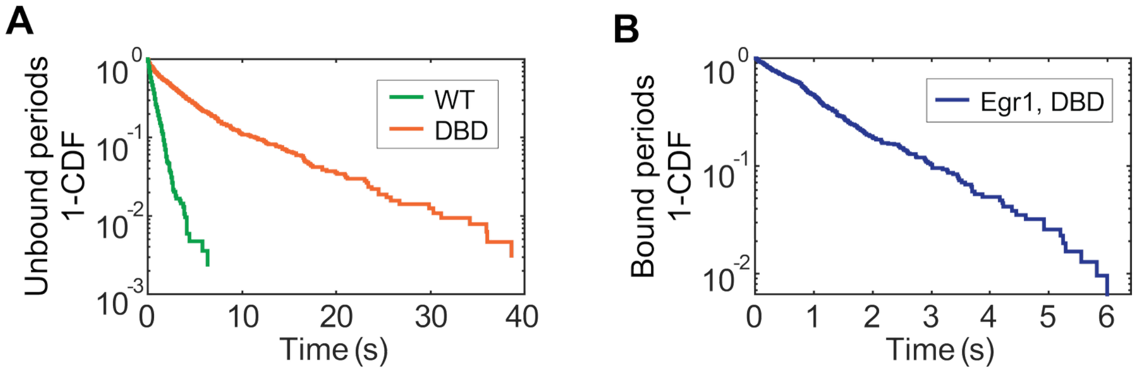


**Figure S3. Supporting results for kinetics experiments**. (A) CDF of the unbound periods, showing a single-exponent distribution for both the WT and DBD variants. (B) CDF of the bound periods for the DBD of Egr1, showing a single-exponential distribution.

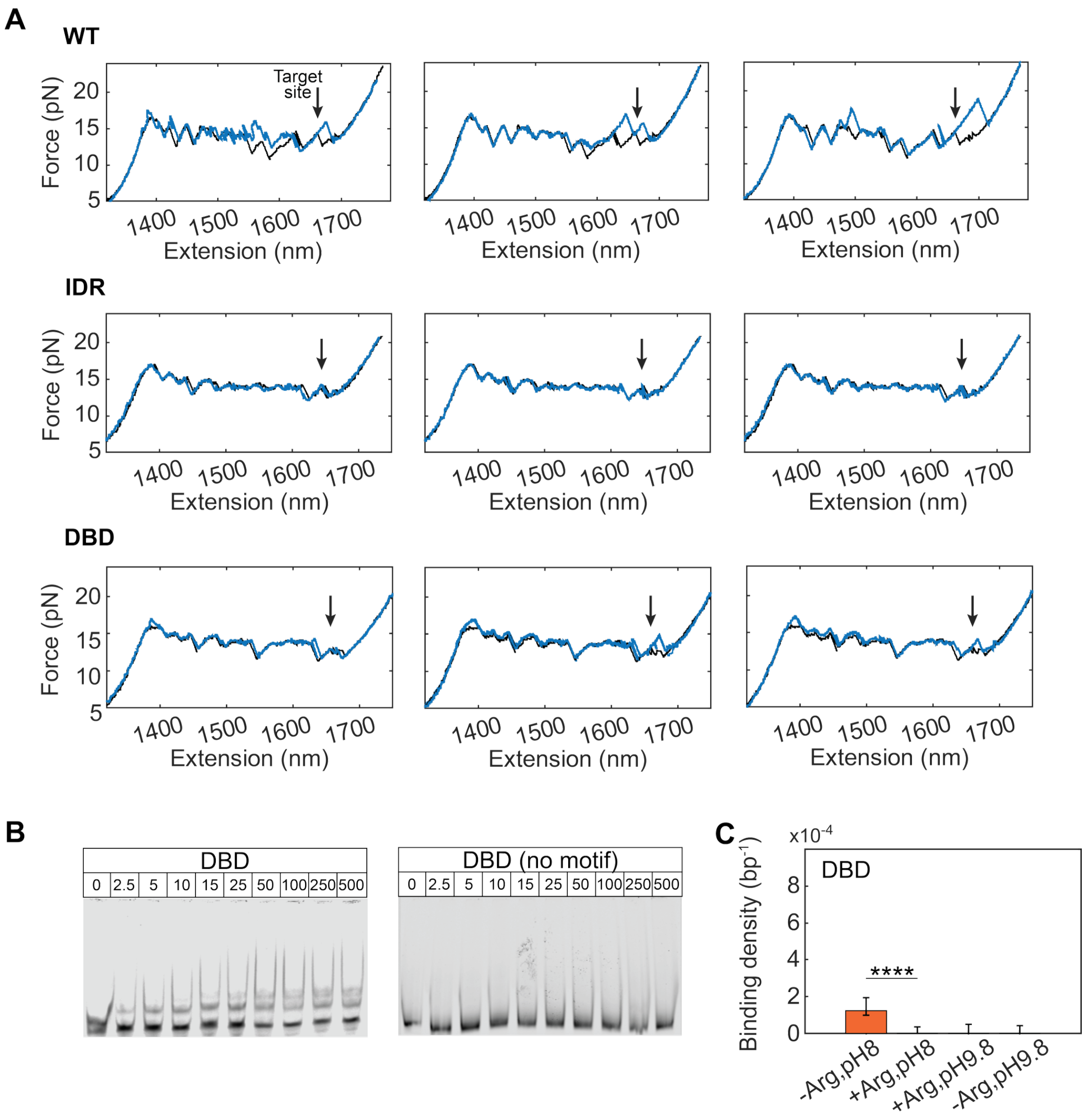
A
WT
Target site
Force (pN)
Extension (nm)
IDR
DBD
B
DBD
DBD (no motif)
0 2.5 5 10 15 25 50 100 250 500
C
x10-4
DBD
Binding density (bp-1)
****
-Arg,pH8
+Arg,pH8
+Arg,pH9.8
-Arg,pH9.8

**Figure S4. Supporting results for Msn2 non-specific binding**. (A) Example of unzipping iteration traces for Msn2 variants, showing various cases of non-specific binding away from the binding site (the binding site is marked by an arrow). For reference, a trace taken at a protein-free solution is shown in black. (B) EMSA using the DBD variant and DNA sequences with and without a binding motif shows no binding in the absence of a binding site. (C) Non-specific binding density of the DBD variant under perturbation conditions as described in Figure 1 (total position bins $n_{-Arg,pH8}$=8959, $n_{+Arg,pH8}$=9734, $n_{+Arg,pH9.8}$=9238, $n_{-Arg,pH9.8}$=10602). Data shown as mean ± SEM, ****P<0.0001, $\chi^2$ test. See also Table S11.

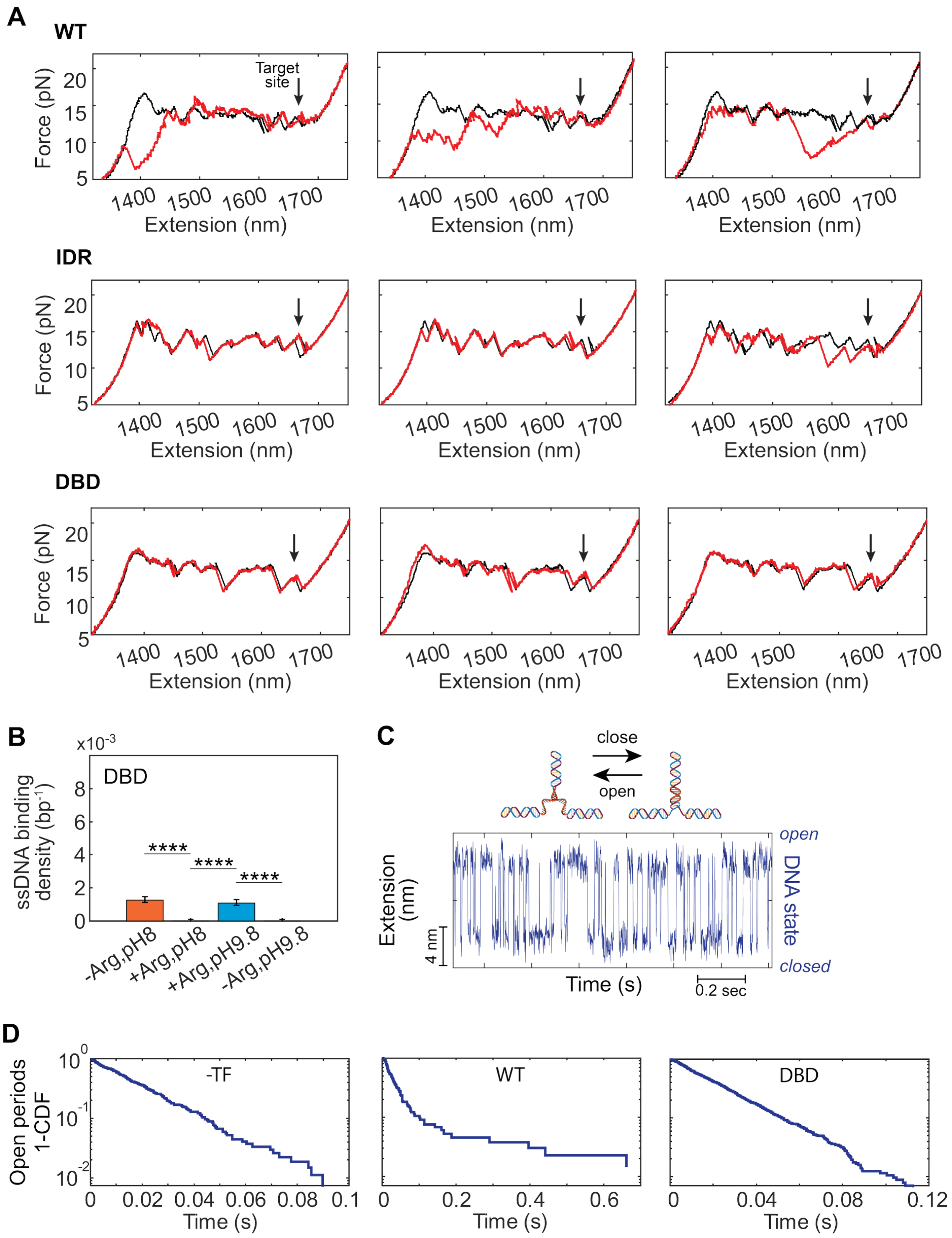
A
WT
Target site
Force (pN)
Extension (nm)
IDR
DBD
B
x10-3
DBD
ssDNA binding density (bp-1)
****
-Arg,pH8
+Arg,pH8
+Arg,pH9.8
-Arg,pH9.8
C
close
open
Extension (nm)
4 nm
open
DNA state
closed
Time (s)
0.2 sec
D
Open periods 1-CDF
-TF
WT
DBD
Time (s)

**Figure S5. Supporting results for Msn2 ssDNA non-specific binding**

(A) Examples of rezipping iteration traces for Msn2 variants, showing various cases of ssDNA non-specific binding (the position of the binding site is marked by an arrow). For a reference, a trace taken at a protein-free solution was added in black. (B) ssDNA non-specific binding density for the DBD variant under perturbation conditions described in Figure 1 (total position bins $n_{-Arg,pH8}=8773$, $n_{+Arg,pH8}=9455$, $n_{+Arg,pH9.8}=6944$, $n_{-Arg,pH9.8}=10230$). Data shown as mean ± SEM, ****P<0.0001, $\chi^2$ test. See also Table S12. (C) Thermal fluctuations between open and closed states during the unbound periods in the kinetics experiment (described in Figure 2). (D) Distribution of open periods with two distinct populations in the presence of the WT variant (but not DBD), indicating ssDNA non-specific binding mediated by Msn2 IDRs.

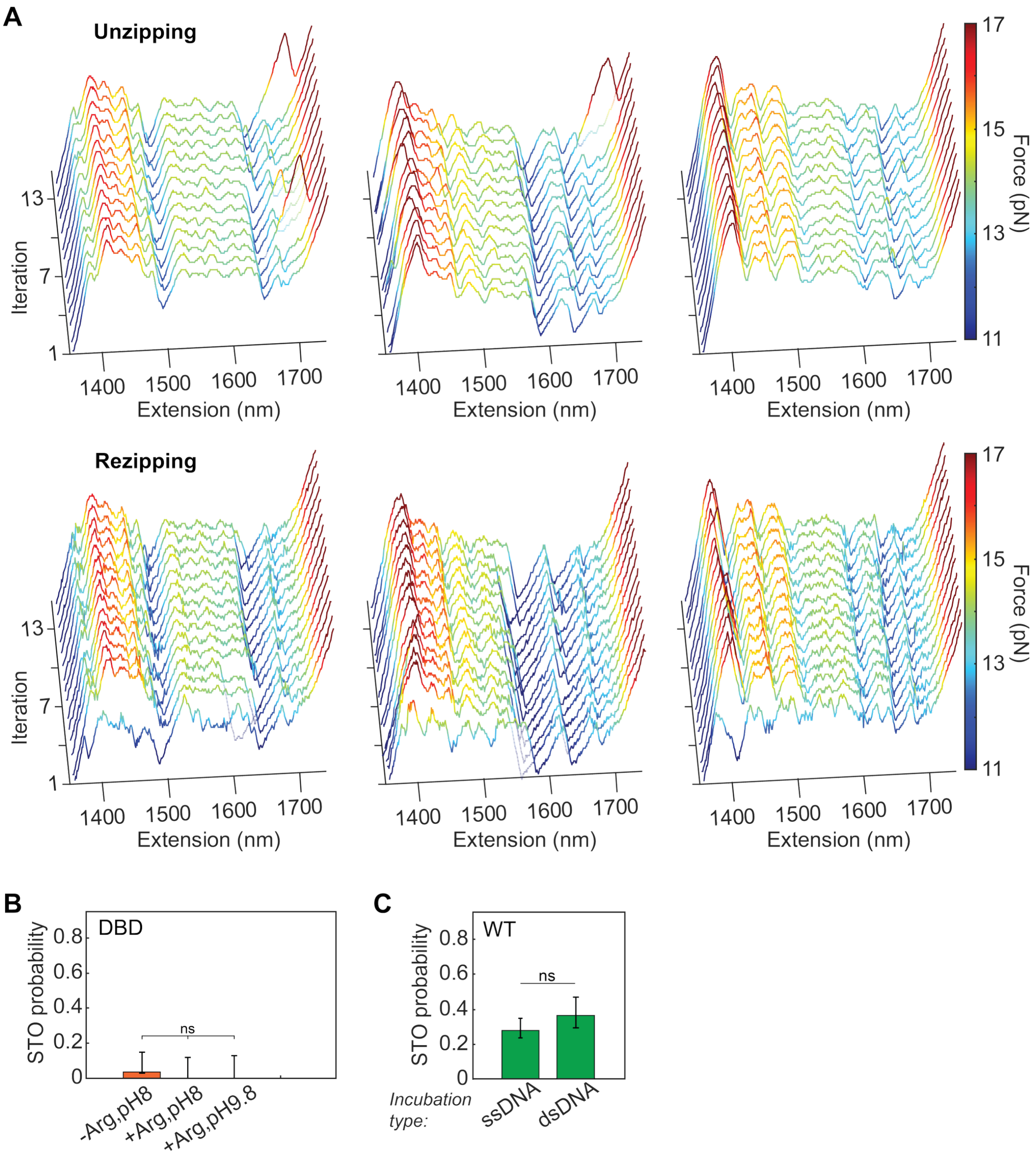


**Figure S6. Supporting results for the IDRs-mediated searching mechanism**

(A) Examples of unzipping and rezipping iterations for WT Msn2 during the detection phase following non-specific incubation, showing various cases of STO binding events. (B) STO probability (as described in Figure 5) for the DBD variant, under electrostatic perturbation conditions as described in Figure 1 (total DNA molecules probed $n_{-Arg,pH8}$=27, $n_{+Arg,pH8}$=24, $n_{+Arg,pH9.8}$=22). Data shown as mean ± SEM, $\chi^2$ test. (C) STO probability for WT Msn2 at unperturbed conditions, following ssDNA and dsDNA incubation (total DNA molecules probed $n_{ssDNA}$=61, $n_{dsDNA}$=28). Data shown as mean ± SEM, $\chi^2$ test. See also Table S13.

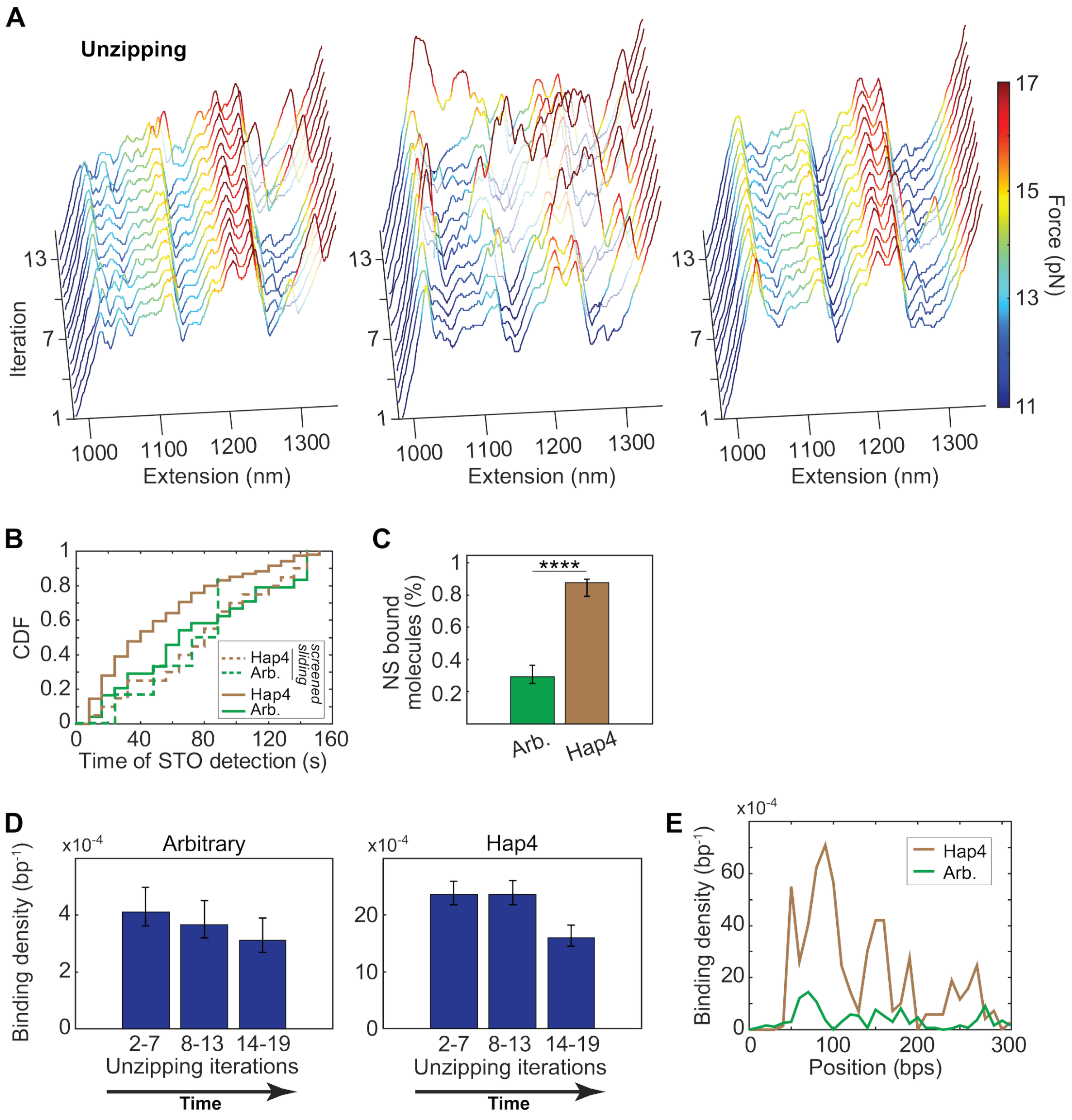


**Figure S7. Supporting results for the sequence dependency of Msn2 searching mechanism**
(A) Examples of unzipping iterations of Msn2 WT for a *Hap4* promoter environment-based DNA construct. (B) Distribution of STO detection times, for *Hap4* and the arbitrary environment, without L-arginine perturbation (full line) and experiments where L-arginine was introduced at the diffusion channel only. (C) Percent of probed molecules that showed any non-specific binding during STO experiments, for *Hap4* and the arbitrary environment (total DNA molecules probed $n_{Arbitrary}$=61, $n_{Hap4}$=41). Data shown as mean ± SEM, ****P<0.0001, $\chi^2$ test. See also Table S14. (D) Non-specific binding density during STO experiments as a function of cycle (and thus time), for the arbitrary (left panel; total position bins $n_{2\text{-}7}$=11253, $n_{8\text{-}13}$=11036, $n_{14\text{-}19}$=11067), and *Hap4* (right panel; $n_{2\text{-}7}$=6789, $n_{8\text{-}13}$=6262, $n_{14\text{-}19}$=6045) environments. The results were clustered into 6-cycles groups. Data shown as mean ± SEM. (E) Non-specific binding density as a function of position on the construct, for *Hap4* and the arbitrary environment.

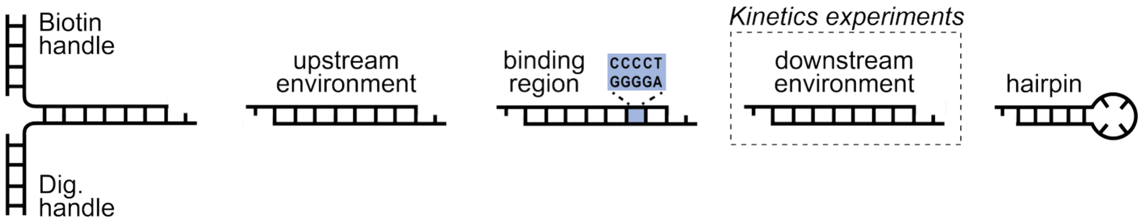


**Figure S8. DNA construct components for the optical tweezers' experiments**

**SUPPLEMENTARY TABLES**

**Table S1. Codon-optimized amino acid sequences of the protein variants**

| Construct | Amino acid sequences |
|---|---|
| WT | MGSSHHHHHHSGSGSAGLEVLFQ\|GPMTVDHDFNSEDILFPIESMSSIQYVENNNPNNINNDVIPYS<br>LDIKNTVLDSADLNDIQNQETSLNLGLPPLSFDSPLPVTETIPSTTDNSLHLKADSNKNRDARTIEN<br>DSEIKSTNNANGSGANQYTTLTSPYPMNDILYNMNNPLQSPSPSSVPQNPTINPPINTASNETNLSP<br>QTSNGNETLISPRAQQHTSIKDNRLSLPNGANSNLFIDTNPNNLNEKLRNQLNSDTNSYSNSISNSN<br>SNSTGNLNSSYFNSLNIDSMLDDYVSSDLLLNDDDDDTNLSRRRFSDVITNQFPSMTNSRNSISHSL<br>DLWNHPKINPSNRNTNLNITTNSTSSSNASPNTTTMNANADSNIAGNPKNNDATIDNELTQILNEY<br>NMNFNDNLGTSTSGKNKSACPSSFDANAMTKINPSQQLQQQLNRVQHKQLTSSHNNSSTNMKSF<br>NSDLYSRRQRASLPIIDDSLSYDLVNKQDEDPKNDMLPNSNLSSSQQFIKPSMILSDNASVIAKVATT<br>GLSNDMPFLTEEGEQNANSTPNFDLSITQMNMAPLSPASSSSTSLATNHFYHHFPQQGHHTMNSKI<br>GSSLRRRKSAVPLMGTVPLTNQQNNISSSSVNSTGNGAGVTKERRPSYRRKSMTPSRRSSVVIESTK<br>ELEEKPFHCHICPKSFKRSEHLKRHVRSVHSNERPFACHICDKKFSRSDNLSQHIKTHKKHGDI |
| IDR | MGSSHHHHHHSGSGSAGLEVLFQ\|GPMTVDHDFNSEDILFPIESMSSIQYVENNNPNNINNDVIPYS<br>LDIKNTVLDSADLNDIQNQETSLNLGLPPLSFDSPLPVTETIPSTTDNSLHLKADSNKNRDARTIEN<br>DSEIKSTNNANGSGANQYTTLTSPYPMNDILYNMNNPLQSPSPSSVPQNPTINPPINTASNETNLSP<br>QTSNGNETLISPRAQQHTSIKDNRLSLPNGANSNLFIDTNPNNLNEKLRNQLNSDTNSYSNSISNSN<br>SNSTGNLNSSYFNSLNIDSMLDDYVSSDLLLNDDDDDTNLSRRRFSDVITNQFPSMTNSRNSISHSL<br>DLWNHPKINPSNRNTNLNITTNSTSSSNASPNTTTMNANADSNIAGNPKNNDATIDNELTQILNEY<br>NMNFNDNLGTSTSGKNKSACPSSFDANAMTKINPSQQLQQQLNRVQHKQLTSSHNNSSTNMKSF<br>NSDLYSRRQRASLPIIDDSLSYDLVNKQDEDPKNDMLPNSNLSSSQQFIKPSMILSDNASVIAKVATT<br>GLSNDMPFLTEEGEQNANSTPNFDLSITQMNMAPLSPASSSSTSLATNHFYHHFPQQGHHTMNSKI<br>GSSLRRRKSAVPLMGTVPLTNQQNNISSSSVNSTGNGAGVTKERRPSYRRKSMTPSRRSSVVIESTK<br>ELDI |
| DBD | MGSSHHHHHHKIEEGKLVIWINGDKGYNGLAEVGKKFEKDTGIKVTVEHPDKLEEKFPQVAATGD<br>GPDIIFWAHDRFGGYAQSGLLAEITPDKAFQDKLYPFTWDAVRYNGKLIAYPIAVEALSLIYNKDLL<br>PNPPKTWEEIPALDKELKAKGKSALMFNLQEPYFTWPLIAADGGYAFKYENGKYDIKDVGVDNAG<br>AKAGLTFLVDLIKNKHMNADTDYSIAEAAFNKGETAMTINGPWAWSNIDTSKVNYGVTVLPTFKG<br>QPSKPFVGVLSAGINAASPNKELAKEFLENYLLTDEGLEAVNKDKPLGAVALKSYEEELAKDPRIAA<br>TMENAQKGEIMPNIPQMSAFWYAVRTAVINAASGRQTVDEALKDAQTTGSGSLEVLFQ\|GPCMEE<br>KPFHCHICPKSFKRSEHLKRHVRSVHSNERPFACHICDKKFSRSDNLSQHIKTHKKHGDI |

Red: N-terminal His6-tag. Cyan: HRV3C protease cleavage site (vertical lines between Q and G indicate cleavage position). Yellow: m
protein (MBP).

**Table S2. Charge of Msn2 WT protein amino acid residues at pH 8.0**

| |
|---|
| MTVDHDFNSE DILFPIESMS SIQYVENNNP NNINNDVIPY SLDIKNTVLD<br>SADLNDIQNQETSLNLGLPPLSFDSPLPVTETIPSTTDNSLHLKADSNKNRDARTIENDSEIKSTNNASGSGANQYTTLTSPYPMNDILYNMNNPL<br>NPPINTASNETNLSPQTSNGNETLISPRAQQHTSIKDNRLSLPNGANSNLFIDTNPNNLNEKLRNQLNSDTNSYSNSISNSNSNSTGNLNSSYFNS<br>LLLNDDDDDTNLSRRRFSDVITNQFPSMTNSRNSISHSLDLWNHPKINPSNRNTNLNITTNSTSSSNASPNTTTMNANADSNIAGNPKNNDATI<br>NFNDNLGTSTSGKNKSACPSSFDANAMTKINPSQQLQQQLNRVQHKQLTSSHNNSSTNMKSFNSDLYSRRQRASLPIIDDSLSYDLVNKQDEDPK<br>QFIKPSMILSDNASVIAKVATTGLSNDMPFLTEEGEQNANSTPNFDLSITQMNMAPLSPASSSSTSLATNHFYHHFPQQGHHTMNSKIGSSLRRRK<br>QQNNISSSSVNSTGNGAGVTKERRPSYRRKSMTPSRRSSVVIESTKE*LEEKPFHCHICPKSFKRSEHLKRHVRSVHSNERPFACHICDKKFSRSDNLS* |

IDR: #1-642 (positively/negatively charged at pH8.0)
*DBD*: #643-704

**Table S3. pKa values of Msn2 IDR residues**

| Amino acid | Side chain $pK_a$ | Isoelectric point | Number of IDR residues |
|---|---|---|---|
| Arginine | 12.5 | 10.8 | 23 |
| Aspartic acid | 3.9 | 3.0 | 43 |
| Glutamic acid | 4.3 | 3.2 | 19 |
| Histidine | 6.0 | 7.6 | 12 |
| Lysine | 10.5 | 9.8 | 22 |

**Table S4. Parameters derived from fitting a two-step binding model to the EMSA data**

| Variant | Binding site | $K_1$ (nM) | $n_1$ | $K_2$ (nM) | $n_2$ | $R^2$ |
|---|---|---|---|---|---|---|
| WT | + | 23.9 ±2.5 | 2.2 ±0.5 | 39.5 ±1.3 | 5.5 ±0.6 | 0.995 |
| | - | 44.5 ±7.5 | 2.1 ±0.5 | 38.3 ±9.1 | 9.1 ±8.2 | 0.999 |
| IDR | + | | | 27.5 ±2 | 2.2 ±0.3 | 0.992 |
| | - | | | 38.7 ±1.7 | 2.3 ±0.2 | 0.997 |

**Table S5. P-values for Figure 1 data**

| 1D<br>Binding probability (Chi-square) | | **WT** |
|---|---|---|
| | **DBD** | <0.0001 |

| 1D<br>Breaking force (t-test) | | **WT** |
|---|---|---|
| | **DBD** | <0.0001 |

| 1F<br>WT, binding probability (Chi-square) | | **-Arg,pH8** | **+Arg,pH8** | **+Arg,pH9.8** |
|---|---|---|---|---|
| | **-Arg,pH9.8** | <0.0001 | <0.0001 | 0.112 |
| | **+Arg,pH9.8** | <0.0001 | <0.0001 | |
| | **+Arg,pH8** | <0.0001 | | |

| 1F<br>DBD, binding probability (Chi-square) | | **-Arg,pH8** | **+Arg,pH8** | **+Arg,pH9.8** |
|---|---|---|---|---|
| | **-Arg,pH9.8** | 0.009 | 0.642 | 0.595 |
| | **+Arg,pH9.8** | 0.038 | 0.327 | |
| | **+Arg,pH8** | 0.003 | | |

**Table S6. P-values for Figure 2 data**

| 2B<br>$k_{on}$ (Permutation test) | | **WT** |
|---|---|---|
| | **DBD** | <0.0001 |

| 2B<br>$k_{off}$ (Permutation test) | | **WT** |
|---|---|---|
| | **DBD** | 0.313 |

| 2D<br>$k_{off1}$, $k_{off2}$ (Permutation test) | | **Koff1, WT** | **Koff1, DBD** | **Koff2, WT** |
|---|---|---|---|---|
| | **Koff2, DBD** | <0.0001 | <0.0001 | <0.0001 |
| | **Koff2, WT** | <0.0001 | <0.0001 | |
| | **Koff1, DBD** | <0.0001 | | |

**Table S7. P-values for Figure 3 data**

| | | | | |
|---|---|---|---|---|
| 3B<br>NS binding density (Chi-square) | | **WT** | **IDR** | |
| | **DBD** | <0.0001 | 0.213 | |
| | **IDR** | 0.032 | | |
| 3H<br>NS binding density (Chi-square) | | **-Arg,pH8** | **+Arg,pH8** | **+Arg,pH9.8** |
| | **-Arg,pH9.8** | 0.076 | <0.0001 | 0.002 |
| | **+Arg,pH9.8** | 0.165 | <0.0001 | |
| | **+Arg,pH8** | <0.0001 | | |

**Table S8 P-values for Figure 4 data**

| | | | | |
|---|---|---|---|---|
| 4B<br>ssDNA NS binding density (Chi-square) | | **WT** | **IDR** | |
| | **DBD** | <0.0001 | 0.779 | |
| | **IDR** | 0.0008 | | |
| 4D<br>ssDNA NS binding density for WT (Chi-square) | | **-Arg,pH8** | **+Arg,pH8** | **+Arg,pH9.8** |
| | **-Arg,pH9.8** | <0.0001 | 0.0004 | <0.0001 |
| | **+Arg,pH9.8** | <0.0001 | 0.373 | |
| | **+Arg,pH8** | <0.0001 | | |

**Table S9. P-values for Figure 5 data**

| | | | | | |
|---|---|---|---|---|---|
| 5C<br>STO probability (Chi-square) | | **-Arg,pH8 (DBD)** | **-Arg,pH8 (WT)** | **+Arg,pH8 (WT)** | **+Arg,pH9.8 (WT)** |
| | **-Arg,pH9.8(WT)** | 0.004 | 0.529 | 0.001 | 0.163 |
| | **+Arg,pH9.8(WT)** | 0.051 | 0.336 | 0.041 | |
| | **+Arg,pH8(WT)** | 0.651 | 0.003 | | |
| | **-Arg,pH8(WT)** | 0.009 | | | |
| 5E<br>STO probability (Chi-square) | | **Arb,None** | **Arb,Sliding** | **Hap4,None** | |
| | **Hap4,Sliding** | 0.241 | 0.464 | <0.0001 | |
| | **Hap4,None** | <0.0001 | <0.0001 | | |
| | **Arb,Sliding** | 0.602 | | | |
| 5F<br>STO binding density (Chi-square) | | **Hap4** | | | |
| | **Arb** | <0.0001 | | | |

**Table S10. P-values for Figure S2 data**

| | | | | |
|---|---|---|---|---|
| S2B<br>WT, binding probability (Chi-square) | | **150KCl,0Arg** | **150KCl,50Arg** | |
| | **200KCl,0Arg** | <0.0001 | 0.959 | |
| | **150KCl,50Arg** | <0.0001 | | |
| S2B<br>DBD, binding probability (Chi-square) | | **150KCl,0Arg** | **150KCl,50Arg** | |
| | **200KCl,0Arg** | <0.0001 | 0.244 | |
| | **150KCl,50Arg** | 0.003 | | |
| S2B<br>WT, Breaking force (t-test) | | **150KCl,0Arg** | **150KCl,50Arg** | |
| | **200KCl,0Arg** | <0.0001 | 0.676 | |
| | **150KCl,50Arg** | <0.0001 | | |
| S2B<br>DBD, Breaking force (t-test) | | **150KCl,0Arg** | **150KCl,50Arg** | |
| | **200KCl,0Arg** | 0.023 | 0.850 | |
| | **150KCl,50Arg** | 0.007 | | |
| S2C<br>WT, Breaking force (t-test) | | **-Arg,pH8** | **+Arg,pH8** | **+Arg,pH9.8** |
| | **-Arg,pH9.8** | 0.224 | <0.0001 | <0.0001 |
| | **+Arg,pH9.8** | <0.0001 | 0.867 | |
| | **+Arg,pH8** | <0.0001 | | |
| S2C<br>DBD, Breaking force (t-test) | | **-Arg,pH8** | **+Arg,pH8** | **+Arg,pH9.8** |
| | **-Arg,pH9.8** | 0.064 | 0.308 | 0.667 |
| | **+Arg,pH9.8** | 0.023 | 0.566 | |
| | **+Arg,pH8** | 0.007 | | |

**Table S11. P-values for Figure S4 data**

| | | | | |
|---|---|---|---|---|
| S4C<br>Binding density of DBD (Chi-square) | | **-Arg,pH8** | **+Arg,pH8** | **+Arg,pH9.8** |
| | **-Arg,pH9.8** | 0.005 | 0.032 | 0.827 |
| | **+Arg,pH9.8** | 0.013 | 0.022 | |
| | **+Arg,pH8** | <0.0001 | | |

**Table S12. P-values for Figure S5 data**

| | | | | |
|---|---|---|---|---|
| S5B<br>ssDNA NS binding density, DBD (Chi-square) | | **-Arg,pH8** | **+Arg,pH8** | **+Arg,pH9.8** |
| | **-Arg,pH9.8** | <0.0001 | 0.463 | <0.0001 |
| | **+Arg,pH9.8** | 0.414 | <0.0001 | |
| | **+Arg,pH8** | <0.0001 | | |

**Table S13. P-values for Figure S6 data**

| | | | |
|---|---|---|---|
| S6B<br>STO probability, DBD (Chi-square) | | **-Arg,pH8** | **+Arg,pH8** |
| | **+Arg,pH9.8** | 0.362 | N/A |
| | **+Arg,pH8** | 0.341 | |
| S6C<br>STO probability, WT (Chi-square) | | **dsDNA** | |
| | **ssDNA** | 0.455 | |

**Table S14. P-values for Figure S7 data**

| | | |
|---|---|---|
| S7C<br>NS bound molecules (Chi-square) | | **Hap4** |
| | **Arb** | <0.0001 |

**Table S15. PCR primers for single-molecule experiment constructs**

| # | Gene promoter | Segment Type | Template | Direction | Sequence (5’-3’) |
|---|---|---|---|---|---|
| 1 | *Cga* | Upstream environment | Mouse genome | Forward | ATCACCACGTGAGACATTTTGAGGTAGTGGTG |
| 2 | | | | Reverse | ATCACGCAGTGCTGTTAATTTAAGAAATTGGAGCAATTGT |
| 3 | *Hap4* | Upstream environment | SC yeast genome | Forward | GCATCACCACGTGTTTTTTTAGAGTAGTCTCGGAAAGA |
| 4 | | | | Reverse | CAGTCACGCAGTGTCCGAAGTCTAAAAGTTTAATCC |
| 5 | *Hor7* | Binding region | SC yeast plasmid | Forward | ATGGCCTTGCTGGCCCTCGAGCACGCTTGTAA |
| 6 | | | | Reverse | ATGGCCTCTATGGCCTGACAATATCTGTATGATTTGATAGT |
| 7 | *Cga* | Downstream environment | Mouse genome | Forward | ATCACTAGGTGTGTTAATTTAAGAAATTGGAGCAATTG |
| 8 | | | | Reverse | ATCACTGCGTGTTATGAAGAGAGAGCATTTGGC |

**Table S16. Annealing oligonucleotides for single-molecule experiment constructs**

| # | Segment | Segment Type | Strand | Sequence (5’-3’) |
|---|---|---|---|---|
| 1 | Msn2 motif + AT-rich sequence | Binding region | Top | ATTTATATAATATAAATTAATATTAATATTGCCCCTGATTATTTAATA TAATTATAAAGCTAG |
| 2 | | | Bottom | GCTTTATAATTATATTAAATAATCAGGGGCAATATTAATATTAATTTA TATTATATAAATGCA |
| 3 | Egr1 motif + 5 flanking bp from *Lhb* | Binding region | Top | AATGTCAGCTAAGCCCTGACACCTGGGCCGAGTGTGAGGCCAATTCAC TCTTGCCACCCCCACAACCCTAG |
| 4 | | | Bottom | GGGTTGTGGGGGTGGCAAGAGTGAATTGGCCTCACACTCGGCCCAGGT GTCAGGGCTTAGCTGACATTGCA |
| 5 | Self-folded hairpin | Hairpin | | /PHOSPHATE/GACTTGAGGCAATTGCCTCAAGTCCTA |
| 6 | Self-folded hairpin (kinetics) | Hairpin | | /PHOSPHATE/GACTTGAGGCAATTGCCTCAAGTCTGC |

**Table S17. Oligonucleotides for EMSA and Mass Photometry experiments**

| # | Segment | | Sequence (5’-3’) |
|---|---|---|---|
| 1 | 51bp DNA w/ motif | top | *CGAAATTATATGGATGAGTAGGTGGGGAAGGAAGTATGTGTAGGTGGCAGG |
| 2 | | bottom | GCTTTAATATACCTACTCATCCACCCCTTCCTTCATACACATCCACCGTCC |
| 3 | 51bp DNA w/o motif | top | *CCTGCCACCTACACATACTTCCTACACAACCTGAGTCGTATTAATTTCGCG |
| 4 | | bottom | CGCGAAATTATACGACTCAAGTTGTGTAGGAAGTATGTGTAGGTGGCGAGG |

* 5′ IRD800 labeling for EMSA experiments
